# Conservation of Regulatory Elements with Highly Diverged Sequences Across Large Evolutionary Distances

**DOI:** 10.1101/2024.05.13.590087

**Authors:** Mai H.Q. Phan, Tobias Zehnder, Fiona Puntieri, Bai-Wei Lo, Boris Lenhard, Ferenc Mueller, Martin Vingron, Daniel M. Ibrahim

**Affiliations:** Berlin Institute of Health at Charité – Universitätsmedizin Berlin, Center for Regenerative Therapies, Charitéplatz 1, 10117 Berlin; Max Planck Institute for Molecular Genetics, Ihnestr. 63, 14195 Berlin; MRC London Institute of Medical Sciences, London, UK; Institute of Clinical Sciences, Faculty of Medicine, Imperial College London, Hammersmith Hospital Campus, London, UK; Institute of Cancer and Genomic Sciences, Birmingham Centre for Genome Biology, College of Medical and Dental Sciences, University of Birmingham, Birmingham, UK

## Abstract

Embryonic gene expression is remarkably conserved across vertebrates as observed, for instance, in the developing hearts of chicken and mouse which diverged >300 million years ago. However, most *cis* regulatory elements (CREs) are highly divergent, which makes orthology tracing based on sequence similarity difficult, especially at larger evolutionary distances. Some evidence suggests functional conservation of CREs despite sequence divergence. However, it remains unclear how widespread such functional conservation might be.

Here, we address this question by profiling the regulatory genome in the embryonic hearts of chicken and mouse at equivalent developmental stages. Gene expression and 3D chromatin structure show remarkable similiarity, while the majority of CREs are non-alignable between the two species. To identify orthologous CREs independent of sequence alignability, we introduce a synteny-based strategy called Interspecies Point Projection (IPP). Compared to alignment-based approaches, IPP identifies up to 5-fold more putative orthologs in chicken, and up to 9-fold across distantly related vertebrates. We term these sequence-diverged orthologs indirectly conserved and characterize their functional conservation compared to sequence-alignable, directly conserved CREs. Indirectly and directly conserved elements show similar enrichment of functional chromatin signatures and cell-type specific enhancer sequence composition. Yet, shared transcription factor binding sites between orthologs are more heavily rearranged in indirectly conserved elements. Finally, we validate functional conservation of indirectly conserved chicken enhancers in mouse using *in vivo* reporter assays. Taken together, by overcoming the limitations of alignment-based methods our results reveal functional conservation of CREs across large evolutionary distances is more widespread than previously recognized.

## Introduction

Embryonic organ development is driven by deeply conserved sets of transcription factors (TF) and signaling molecules that control tissue patterning, cell fates, and ultimately morphogenesis. Especially during the phylotypic stage, but also later during organogenesis many lineage and tissue-specific gene expression patterns are similar even between distantly related organisms (Irie and Kuratani 2011; Berthelot et al. 2017). One such example is the developing heart, where cellular patterning and morphological changes are deeply conserved across vertebrate lineages. The same key group of TFs expressed in the cardiac mesoderm is required for organogenesis, from the two-chambered heart in fish to the four-chambered hearts of birds and mammals (Olson 2006). Thus, the TFs of this gene regulatory network argue for a common genetic basis of embryonic development. Moreover, coding mutations in these genes have been shown account for ∼45% of congenital heart disease (CHD) cases, the most common human birth defect (Pediatric Cardiac Genomics Consortium et al. 2013; Zaidi et al. 2013; Jin et al. 2017). Many of the ∼55% unsolved cases, not only for CHD, but also for other genetic diseases, might be caused by non-coding variants perturbing CREs of those developmental genes (Richter et al. 2020; Xiao et al. 2024).

However, many cis-regulatory elements (CREs) detected experimentally through DNA-accessibility or chromatin modifications are not sequence conserved (Visel et al. 2009; Villar et al. 2015), especially across larger evolutionary distances. For example, enhancers identified by chromatin modifications in embryonic heart tissues are poorly conserved (Blow et al. 2010). Similar observations were shown for TF binding sites (TFBS) in livers of five different vertebrate species (Schmidt et al. 2010). Yet, there are several examples for functionally conserved CREs in the absence of sequence conservation (Fisher et al. 2006; McGaughey et al. 2008; Madgwick et al. 2019). For example, the well-known *even-skipped* stripe 2 enhancer shows functional conservation amongst insects despite highly divergent sequences (Ludwig, Patel, and Kreitman 1998; Hare et al. 2008; Crocker and Stern 2017).

Determining orthologous CREs in distantly related species is complicated for several reasons. One, the rapid turnover in non-coding sequences constrains the effectiveness of pairwise alignments. Two, alignment-free methods struggle to accurately determine ortholog pairs. Alignment-free methods search for similar clusters of TFBS or “sequence words” as footprints of regulatory elements. (Sanges et al. 2006; Vinga 2014; Zielezinski et al. 2017, 2019). A more recent alignment-free approach, machine-learning algorithms, were successfully employed to identify cell-type specific enhancers across species. While this highlights the conservation of regulatory information independent of sequence alignability (Minnoye et al. 2020; Oh and Beer 2023; Kaplow et al. 2023; Kliesmete et al. 2024), additional processing steps are needed to establish putative ortholog pairs. Three, the computational demands and availability of genome assemblies limit the use of multiple genome alignments, which is an alternative better suited to the task of orthology tracing across species. For example, the zebrafish ortholog of a human limb enhancer was identified indirectly through iterative pairwise alignment between human and spotted gar, and between spotted gar and zebrafish (Braasch et al. 2016). More systematically, the use of one bridging species (Xenopus) helped to uncover hundreds of such “covert” ortholog pairs between human and zebrafish (Taher et al. 2011). Using Cactus multi-species alignments from hundreds of genomes, approaches like halliftover/HALPER (Paten et al. 2011; Hickey et al. 2013; Zhang et al. 2020; Armstrong et al. 2020) aim to trace orthology from genome sequences alone. However, in addition to the required computational infrastructure and the availability of genome assemblies, these approaches are currently not available for larger evolutionary distances (e.g. chicken-mouse).

Here we present an experimental-analytical framework to efficiently identify orthologous CREs combining two currently underutilized features – synteny and functional genomic data. In genomics, *synteny* describes the maintenance of colinear genomic sequences on chromosomes of different species (Engström et al. 2007; Kikuta et al. 2007). Not only genes are maintained in synteny; developmental genes are often flanked by conserved non-coding elements (CNEs), many of which act as enhancers (Siepel et al. 2005; Bejerano et al. 2005; Visel, Bristow, and Pennacchio 2007). Their syntenic arrangement reflects conserved regulatory environments that have been described as genomic regulatory blocks (GRBs)(Kikuta et al. 2007; Harmston et al. 2017). *Functional genomic data*, such as chromatin accessibility and histone modifications, are widely used to determine putative CREs in any tissue of interest. Given that the hearts of birds and mammals are evolutionary homologous structures, the active regulatory genome in both should be related. Therefore, experimentally identified CREs in both species might provide the genomic footprint of functionally conserved orthologs whose sequences have diverged to the point where alignment fails. We first use chromatin profiling from murine and chicken hearts at equivalent developmental stages to experimentally determine regulatory elements. We then apply Interspecies Point Projection (IPP), a synteny-based algorithm designed to map corresponding genomic locations in highly diverged genomes.

Using this strategy, we uncover thousands of previously hidden orthologous CREs based on their relative position in the genome and overcome current limitations. We term these sequence-diverged orthologs indirectly conserved, and validate their functional equivalence as compared to classical sequence-conserved elements. We find similar enrichment of chromatin marks at directly and indirectly conserved elements. Furthermore, using machine learning models and TFBS-driven analysis, we show that both classes display similar heart-enhancer specific sequence composition. Yet, shared TFBS are more heavily rearranged between indirectly conserved CRE pairs. Finally, we demonstrate their functional orthology using *in vivo* enhancer-reporter assays. Thereby we demonstrate a currently underrepresented widespread conservation of cis-regulatory elements with highly diverged sequences across large evolutionary distances.

## Results

### Identification of heart CREs from equivalent developmental stages in mouse and chicken

To identify the cis-regulatory elements driving gene expression at equivalent stages of heart development, we generated comprehensive chromatin and gene expression profiles from embryonic mouse and chicken hearts (ChIPmentation, ATAC-seq, RNA-seq, Hi-C) at E10.5/E11.5 and HH22/HH24 (**Fig. 1a**). To compare global gene expression profiles, we measured differentially expressed genes in the heart vs. limb in both mouse and chicken (**Fig. 1b**). Consistent with previous reports (Olson 2006) tissue-specific expression is conserved, including key TF genes specific for heart and limb development (**Fig. 1b, Fig. S1a**). To characterize conservation of regulatory regions driving this expression, we first estimated sequence conservation by alignment of open chromatin regions using LiftOver (Kuhn et al. 2009). Most mouse peaks in non-coding regions lacked sequence conservation in chicken, in stark contrast to those overlapping exons (**Fig. 1c, Fig. S1b**). We then used Hi-C and ChIPmentation data to more comprehensively profile the regulatory genome. Hi-C showed global conservation of the 3D genome in syntenic regions surrounding most developmental genes (**Fig. 1d**) and enrichment of synteny breaks at TAD boundaries (**Fig. S1**). Syntenic regions surrounding developmental genes showed comparable distribution of chromatin marks indicating that the position of regulatory elements relative to their targets might be conserved (**Fig. 1d**). We used CRUP to predict active CREs from typical histone modifications (Ramisch et al. 2019). To further refine our list, we integrated CRUP predictions with chromatin accessibility and gene expression data, followed by stringent filtering (see Methods) to have a high-confidence set of active enhancers and promoters for both species, minimizing the number of false-positive regions. In total, we called 20,252 promoters and 29,498 enhancers in mouse and 14,806 and 21,641 in chicken hearts, respectively (**Fig. 1d, Fig. S1c**).

**Figure 1:**
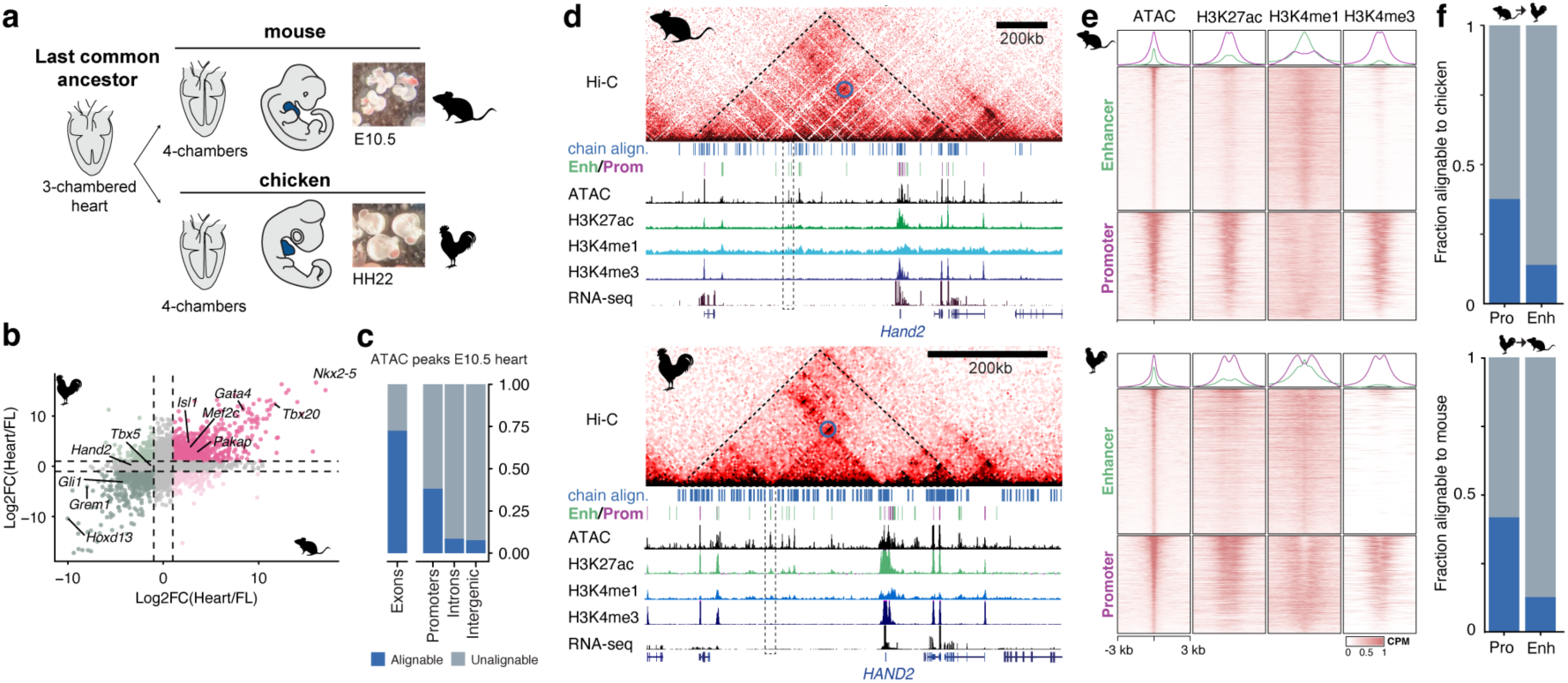
Evolutionary conservation of gene expression and chromatin structure between mouse and chicken embryonic hearts despite divergent cis-regulatory elements. a) Reptilian and mammalian lineages convergently evolved fully separated 4-chambered hearts. E10.5/HH22 represent equivalent stages of heart formation b) Conservation of global gene expression (log2 fold-change of heart vs. limb expressed genes) between mouse (E10.5) and chicken (HH22). c) ATAC-seq peaks (E10.5 heart) are mostly alignable (LiftOver (--minMatch = 0.1)) to chicken in coding, but not in non-coding regions. d) Syntenic regions at the *Hand2/HAND2* locus shows conserved 3D chromatin structure and histone modifications relative to the target gene despite different genomic size. Dashed triangle outline conserved TAD structure, blue circles/dashed rectangle show specific contacts to conserved enhancers. Blue ticks: conserved sequences, Green/Purple ticks: predicted promters/enhancers. e) Signal enrichment (+/-3kb) of histone modifications at heart promoters and enhancers, centred on ATAC-seq peaks f) Fraction of alignable elements identified in e) with the chicken/mouse genome

We then estimated the degree of sequence conservation for this high-confidence set of regulatory elements. Consistent with previous reports (Blow et al. 2010) less than 50% of promoters and only ∼10% of enhancers were sequence-conserved between mouse and chicken (**Fig. 1f, Fig. S1d**). Thus, the lack of sequence alignablity remains consistent, even when restricting the analysis to a stringently filtered set of enhancers and promoters, contrasting conserved gene expression patterns and 3D chromatin structure.

### A synteny-based strategy to identify orthologous genomic regions

Because enhancer function can be maintained despite rapid turnover of underlying sequences, DNA sequence conservation alone likely underestimates conserved regulatory activity. To identify such conserved, non-alignable CREs we developed a synteny-based algorithm, Interspecies Point Projection (IPP) (Baranasic et al. 2022), designed to find orthologous regions independent of sequence divergence (see Supplemental Text and Fig. S2). The approach is based on conserved synteny. We assume any non-alignable element in one genome located between flanking blocks of alignable regions is located at the same relative position in another genome (**Fig. 2a**). Thus, for a given species pair we can interpolate the position of an element (e.g. an enhancer) relative to adjacent alignable regions, so-called *anchor points*. We refer to the interpolated coordinates in the target genome as *projections*. Because a larger distance to an *anchor point* reduces accuracy of projections, the second pillar of IPP involves optimizing the use of bridged/tunneled alignments (Taher et al. 2011; Braasch et al. 2016). IPP uses not one, but multiple bridging species, which increases the number of anchor points thereby minimizing this distance (**Fig. 2b**). With this, IPP classifies orthologous regions by their distance to a *bridged alignment* or *direct alignment*. Regions projected within 300bp to a *direct alignment* are defined as *directly conserved* (DC). Those further than 300bp to a *direct alignment* but which can be projected through bridged alignments we define as *indirectly conserved* (IC) regions if the summed distance to anchor points is less than 2.5kb. The remaining projections are defined as non-conserved (NC) (see **Fig. S2** and Supplemental Text for details and parameterization).

**Figure 2:**
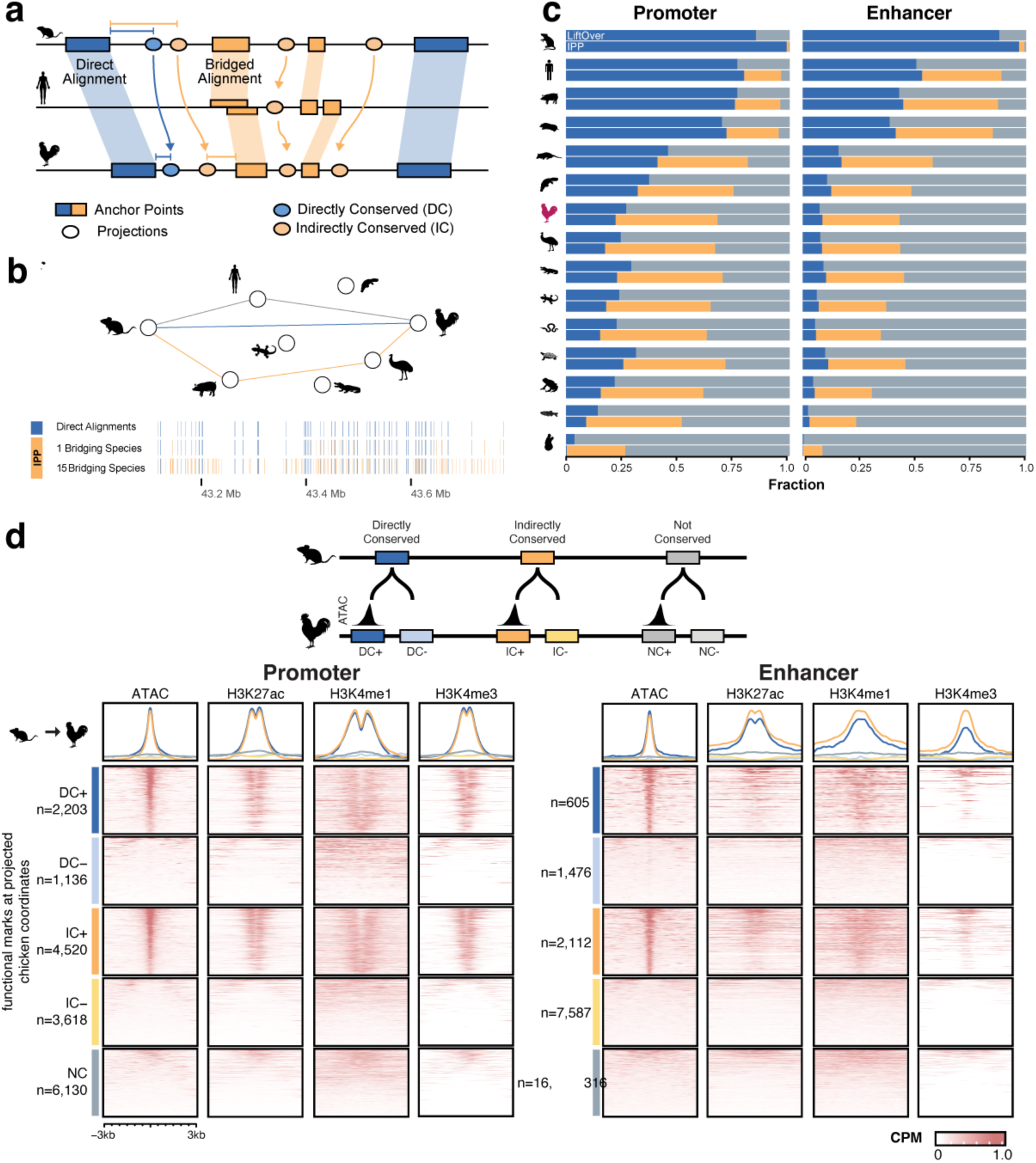
A synteny-based algorithm, Interspecies Point Projection (IPP), identifies thousands of putative sequence orthologs of mouse heart CREs with functional chromatin signatures in chicken. a) Synteny-based proximity to direct/indirectly aligned regions determines orthology between features (e.g. ATAC-peak summits). b) Multi-species bridged alignments increase the number of anchor points in a representative region using 0, 1 and 15 bridging species c) IPP increases the number of putatively homologous regions from mouse to 15 other species used as bridging species (compare blue vs. orange portion). LiftOver alignments (top bar) are compared to IPP *Directly Conserved* and *Indirectly Conserved*. Increase is particularly high at greater evolutionary distances to non-mammalian species d) Classification of elements with or without conserved activity +/-. Signal enrichment at chicken genomic regions to which mouse E10.5 heart elements were projected.

### IPP improves detection of orthologs between distantly related species

To optimize our mouse-chicken projections, we selected a set of 16 species, consisting of mouse, chicken and 14 bridging species from the reptilian and mammalian lineages along with additional ancestral vertebrate/chordate genomes (see Methods). After building our collection of anchor points from pairwise alignments, we project our set of murine heart CREs to chicken and all bridging species to estimate their conservation at varying evolutionary distances. In parallel we used UCSC LiftOver to serve as a reference for sequence conservation for IPP projections. In practice, LiftOver performed similarly to IPP DC projections for all 15 species (**Fig. 2c**), with the exception that multiple mappings can occur when ‘lifting’ the entire sequence. The proportion of mouse CREs classified as *directly conserved* (DC) reduces drastically with increasing evolutionary distances. While over 90% of CREs are conserved when comparing mouse to the closely related rat, this number drops to 50-70% within placental mammals and even more so to non-mammalian vertebrates. Specifically for chicken, only 22% of all promoters and 10% of enhancers are sequence conserved (**Fig. 2c**).

By additionally identifying indirectly conserved (IC) regions, IPP increases the number of conserved elements in all species. Especially within distantly related vertebrates this increases by a factor of 3 to 9, and substantially adds to the number of putatively conserved CREs (orange fraction, **Fig. 2c**). For the mouse-chicken comparison, the percentage of conserved promoters increases 3-fold (18,9%, DC) to 65%, DC+IC), and for enhancers 5-fold (7,4% to 42%). With this, IPP pairs an additional 8,138 and 9,699 promoters and enhancers with candidate ortholog sequences in chicken.

Unlike the synteny-based approach of IPP, other efforts to improve ortholog identification include hierarchical alignments, which are multiple-genome alignments guided by evolutionary relationships (Hickey et al. 2013; Zhang et al. 2020). We compared IPP with halliftover/HALPER (Zhang et al. 2020), which uses Cactus alignments from hundreds of mammalian genomes, for all placental mammals in our species collection (i.e. rat, human, pig, mole). Depending on parameterization, IPP performs similarly or better at identifying orthologous enhancers within this relatively short evolutionary distance. This indicates that ortholog can be traced across evolutionary distances by comparing hundreds of genome sequences. However, the synteny-based strategy of IPP achieves comparable detection rates using only 16 species and spans a greater evolutionary distance than currently available for hierarchical alignments.

Since IPP can project any set of genomic coordinates, we next used IPP on a set of limb enhancers we identified, as well as on two published datasets that reported low conservation between mouse and chicken: murine heart enhancers from (Blow et al. 2010), and a set of CEBP/A TFBSs in liver from (Schmidt et al. 2010). IPP increased the number of putative ortholog heart enhancers equivalent to that of our heart enhancers (**Fig. S2b**). Heart enhancers were slightly less well conserved (DC and IC) than limb enhancers (**Fig. S2c**), confirming general trends observed previously (Blow et al. 2010). For CEBP/A only 2% of murine peaks were reported to be conserved in chicken, and even less bound by CEBP/A in chicken livers (Schmidt et al. 2010). We re-analyzed the ChIP-seq data from mouse and chicken livers and confirmed that only a small fraction (5,7%) of mouse CEBP/A binding sites were directly conserved in chicken (DC) and just 173 of these sites overlapped with a CEBP/A peak in chicken (**Fig. S2f**). However, by including IC projections, we increased the number conserved CEBP/A sites to 32% and found an additional set of 579 peaks that were also CEBP/A bound in chicken livers.

Taken together, IPP dramatically increases detection of orthologous genomic regions, particularly for larger evolutionary distances, uncovering a previously hidden set of conserved elements that can be investigated for their role in evolution and gene regulation.

### Indirectly and directly conserved CREs show a similar enrichment for functional chromatin marks

The large additional number of IC regions suggests that up to 80% of conserved CREs might have gone undetected in most analyses to date. Since we collected functional genomic data from developmentally equivalent stages, we first profiled chromatin signal and compared how well the chromatin state at DC and IC predicted CREs is conserved in chicken. For DC CREs, we found that 66% mouse promoter and 29% enhancer projections overlapped an ATAC-seq peak in the chicken genome. Interestingly, these percentages were similar for IC CREs with 56% promoter and 26% enhancer projections, although the absolute numbers of IC promoters and enhancers is substantially higher than DC. We classified these regions with conserved activity as DC+/IC+ and those without an ATAC-seq peak at the projected site as DC-/IC- (**Fig. 2d**). Consistent with the ATAC-seq signal, DC+/IC+ CREs showed equivalent specific enrichment of H3K4me3 at promoters and H3K4me1 at enhancers, suggesting that the IC CREs identify the functional orthologs of murine heart CREs in the chicken genome (**Fig. 2d**). This similar enrichment of functional chromatin marks suggests that interpolated regions point to “functionally conserved” CREs in the target genome and that sequence homology is an incomplete indicator of conserved activity.

### SVM model robustly learns tissue-specific sequence features and independently validates IPP performance

Recently, machine learning (ML) methods have become a viable strategy to identify cell-type specific CREs in distantly related species, by virtue of their ability to capture complex sequence-function relationships without relying on strict sequence conservation (Minnoye et al. 2020; Oh and Beer 2023; Kliesmete et al. 2024; Kaplow et al. 2023). To test the regulatory content in the IPP projections in chicken, we first trained a gapped k-mer Support Vector Machine (gkm-SVM) model on mouse data to identify heart-specific enhancers. To learn predictive heart-specific enhancer vocabularies, we trained the SVM on aggregated tissue-specific ATAC-seq peaks from mouse embryonic heart outside promoter regions, against the background of non-overlapping peaks from non-heart cell/tissues (**Fig. 3a**, see Methods).

**Figure 3:**
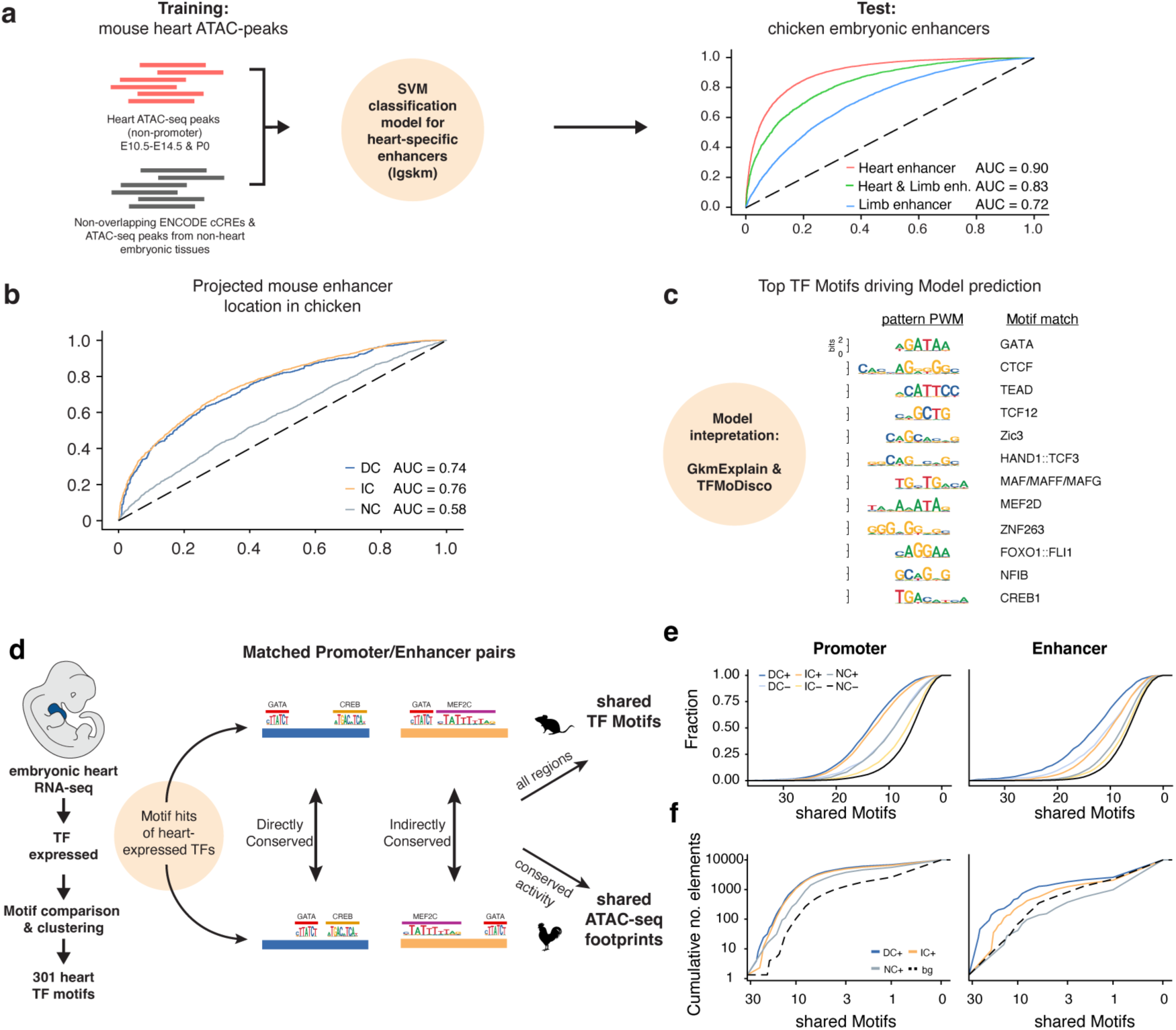
In silico analysis of sequence composition and motif content of indirectly and directly conserved elements. a) Training of a Support Vector Machine (SVM) model to identify heart enhancers with independent data from public repositories. Positive set: embryonic heart/cardiomyocyte ATAC-seq peaks, Negative Set: non-overlapping ATAC-seq peaks from non-heart tissues. The model distinguishes heart- vs. limb-specific enhancers from chicken embryos. AUC: Area Under Curve b) Evaluation of DC+, IC+ and NC classified regions of the chicken genome by the SVM Model. c) TFMoDisco interpretation of the putative TFBS that contribute to model specificity. BS of several known heart-specific TFs contribute to model accuracy. d) Heart-expressed TFs identified from RNA-seq were consolidated to 301 motifs of heart specific TFs. Promoter/Enhancer pairs were screened for shared TFBS or ATAC-seq footprints e) DC+/IC+ promoters/enhancers share more heart TFBS than DC-/IC-, or non-conserved NC regions f) Functionally conserved DC and IC ATAC-seq peak pairs share more TF-footprints than NC ATAC-seq peak pairs or control pairs (bg = a non-paired ATAC-seq peak in the same TAD)

We then tested the model’s cross-species predictive power on the chicken enhancer regions we identified in the embryonic heart and forelimb (FL). The mouse-trained SVM correctly distinguished between heart-specific, shared, and FL-specific chicken enhancers (**Fig. 3a**). This validated that the features from mouse sequences are in fact predictive of heart-specific enhancers in chicken. A recent study found that tissue-specific CREs show a lower degree of sequence conservation than more pleiotropic CREs (Kliesmete et al. 2024). We therefore evaluated SVM-predicted tissue-specificity of all ATAC-Seq peaks from chicken embryonic hearts and noted a clear inverse relationship to sequence conservation (**Fig. S3**). In other words, predicted heart-specific chicken regions (i.e. positive score) are more sequence-divergent from mouse than more pleiotropic peaks, providing further evidence that sequence alignability is a poor estimator of conserved regulatory activity.

Since IC elements exhibit similar degree of conserved activity to DC elements in terms of epigenomic signatures (**Fig. 2d**), we next wanted to estimate conservation as defined by its shared tissue specificity between species. We therefore compared the predicted tissue-specificity of mouse enhancers projected to orthologous chicken loci between DC, IC and NC elements. DC and IC projections were equally likely to be classified as heart-specific enhancers (AUC, DC=0.74, IC=0.76). NC projections, however, were less likely to be classified as heart enhancer (AUC=0.58) (**Fig. 3b**), further indicating conserved tissue-specific enhancer activity.

To better understand predictive sequence patterns learned by the model, we computed the contribution of individual nucleotides from input sequences to the SVM output classification with GkmExplain (Shrikumar, Prakash, and Kundaje 2019) and consolidated recurring high scoring patterns, or ‘seqlets’, into motifs (Shrikumar et al. 2018). Motifs discovered from mouse and chicken sequences largely overlap (**Fig. S3**), suggesting conserved enhancer vocabularies. In fact, known motifs of master regulators of heart development (e.g. GATA, TEAD and HAND) were most predictive of tissue specificity (**Fig. 3c**), further supporting the model’s robustness in predicting heart-specific enhancers. Thus, this independent approach validates that the IPP projections of mouse enhancers faithfully identify heart-specific enhancer regions in the chicken genome.

### Transcription factor binding site conservation as indicator of conserved CRE activity

If IPP projections represent conserved pairs of CREs, these regions should share the same TFBS. Here, we can evaluate this both at the sequence- and chromatin level given our available data using TF motif scanning and ATAC-seq footprinting. We used our heart RNA-seq data to identify TFs expressed in the heart and curated a set of 301 heart TF motifs (**Fig. 3d**). We then calculated for every mouse-chicken ortholog pair how many TFBS were shared and plotted the results (**Fig. 3e**).

Overall, orthologous promoter regions shared more TFBS hits than enhancers. DC+/IC+ promoters were comparable in the number of shared TFBS and both were clearly distinguishable from DC-/IC- promoters (**Fig. 3e**). For enhancers, DC+ shared the most TFBSs, while IC+ enhancer pairs shared as many TFBS as DC-enhancers. Notably, CREs with conserved active chromatin marks (dark blue/orange lines) in all comparisons shared more TFBS than those without (light blue/orange lines), irrespective of direct or indirect conservation. This suggest that functionally conserved orthologs are more likely to retain regulatory information. Finally, we used our ATAC-seq data to compare shared TF footprints. We compared all DC/IC/NC pairs that had ATAC-seq signal in both genomes relative to background (non-orthologous ATAC-seq peaks within the same TAD, see Methods). Consistent with our TFBS motif results, all projection pairs outperformed control regions. DC and IC promoters were equal in the number of shared TF-footprints, while DC enhancers were overall slightly more likely to share TF-footprints than IC enhancers (**Fig. 4f**). These results confirm that IPP identifies orthologous pairs of CREs with shared TFBS, representing a conserved sequence syntax that is independent of direct sequence conservation.

**Figure 4:**
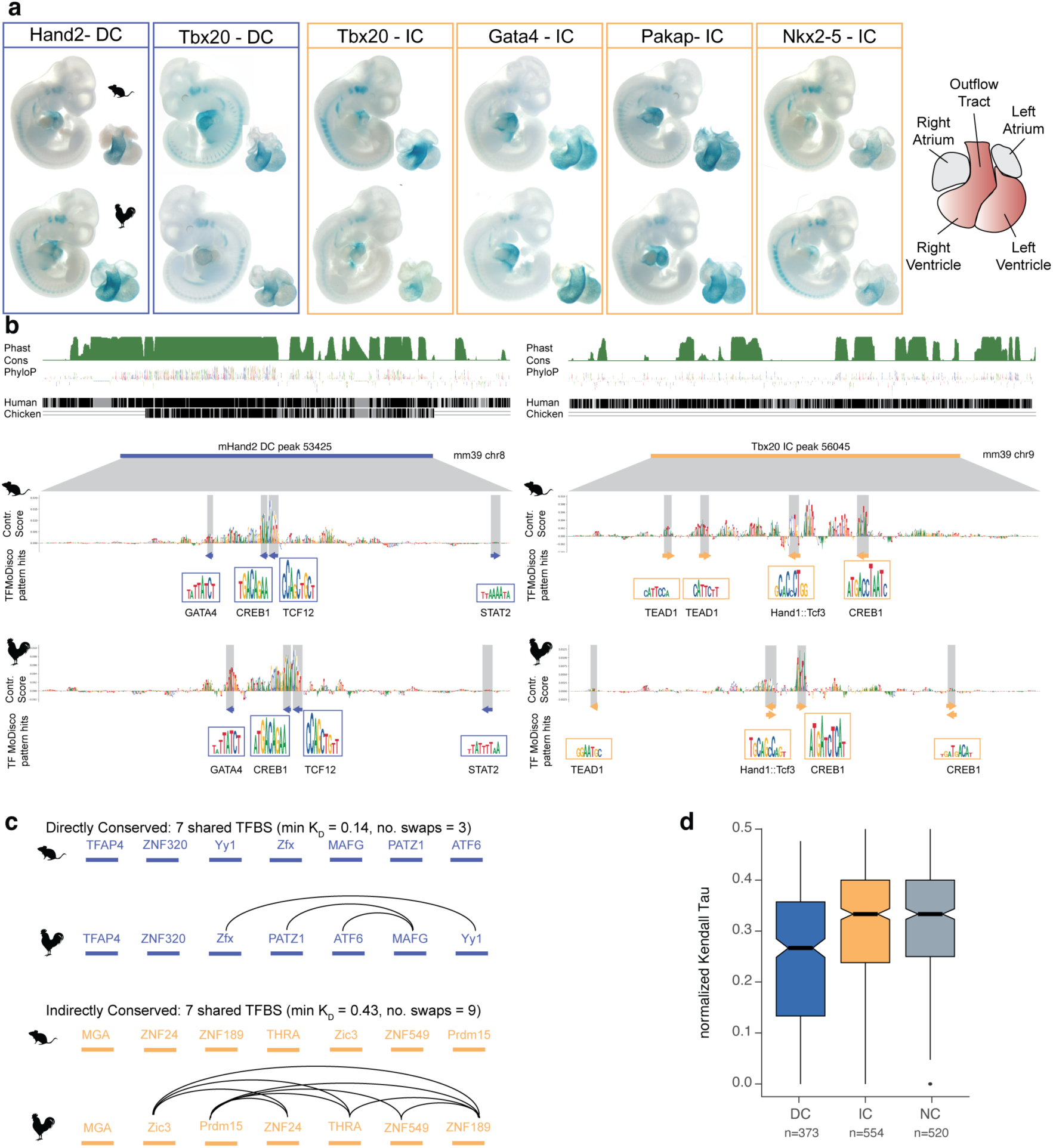
Indirectly conserved heart enhancers from mouse and chicken drive conserved gene expression pattern *in vivo*. a) Directly and Indirectly conserved enhancers from mouse (top) and chicken (bottom) drive highly similar expression patterns in the heart of E10.5 embryos. Individual enhancer show similar tissue-restricted or broad expression patterns. b) Sequence conservation scores (PhastCons/PhyloP) and direct alignments to human and chicken of the murine Hand2-DC and Tbx20-IC enhancer tested in a). SVM contribution scores and TF-Modisco Motif matches show conserved sequence features of the 500bp enhancer highlighting shared TF-Motif hits overlapping with seqlets. c) The different order of shared TFBSs in IC and DC enhancer pairs is reflected in the computed Kendall-Tau Distance, K_D_. d) K_D_ scores for all functionally conserved DC/IC/NC CRE enhancer pairs. Asterisks indicate the magnitude of effect size based on Cohen’s *d*: small (*, *d* < 0.2), medium (**, d ≤ 0.5)

### Indirectly conserved heart enhancers from chicken drive conserved gene expression patterns in mouse embryonic hearts

Gene regulatory elements drive tissue and cell-type specific expression. Based on our analysis, directly and indirectly conserved elements are functionally conserved orthologs and should drive conserved expression patters in the developing heart. To test this, we selected two pairs of DC and 4 pairs of IC enhancers and generated *in vivo* enhancer-reporter mice for each of these elements. We profiled enhancer activity using lacZ staining in E10.5 mouse embryos. All enhancer pairs drove conserved expression with remarkable specificity (**Fig. 4a**). Enhancers driving expression patterns in specific regions of the heart, such as the outflow tract and atrio-ventricular canal, were consistent with those from nearby genes (Hand2-DC, Tbx20-IC, Nkx2-5-IC) (Overbeek 1997; Srivastava and Olson 1997; Firulli et al. 1998; McFadden et al. 2000; Stennard et al. 2003; Prall et al. 2007). The same was true for the ventricle-specific expression of two other enhancers (Tbx20-DC, Gata4-IC) (Heikinheimo, Scandrett, and Wilson 1994). An indirectly conserved enhancer at the *Pakap* locus (Pakap-IC), which contains the *A-kinase anchoring protein 2* (*Akap2*) gene involved in general cardiomyocyte function (Maric et al. 2021), drives broad expression in all cardiac tissues. We integrated scores obtained from the SVM model for all tested pairs. Many seqlets with high contribution scores to our enhancer prediction overlapped with predicted binding sites of key TFs (**Fig. 4b** shaded boxes and **Fig. S4**) and were shared between mouse and chicken CREs for each pair. These data show that the chicken IC enhancers we identify constitute *bona fide* orthologs to their mouse counterparts, regardless of sequence conservation.

### Indirectly conserved CREs show a higher degree of TFBS shuffling

In all our analyses and validations, IC regions showed similar signatures of functional conservation to DC, despite lack of alignablity. We therefore wanted to explore how the underlying DNA sequences may differ in ways they encode regulatory information. We hypothesized that for CRE pairs with a similar number of shared TFBSs, DC pairs would display a more conserved TFBS order within the element than IC pairs (**Fig. 4c**). For example, a DC and IC enhancer pair with both 7 shared TFBSs, show a more shuffled order between IC pairs, likely complicating alignment of the two sequences. To systematically evaluate this phenomenon, we calculated the Kendall-Tau rank distance for all enhancer pairs. The Kendall-Tau rank distance assesses the similarity between two ranked lists by measuring the number of swaps needed to change one list into the order of the other list (Qian and Yu 2019). We selected all functionally conserved enhancer pairs with at least 6 shared TFBSs and computed the normalized Kendall-Tau Distance for each pair (**Fig. 4 c,d**). DC enhancers exhibited a significantly lower KD score (median = 0.27) than IC (median=0.33) and NC enhancers (median=0.33). Consequently, conservation of an element’s regulatory function is likely less dependent on exact sequence conservation than on preserving the appropriate balance of TFBSs within the given element.

## Discussion

Here, we show widespread conservation of functional gene regulatory elements in the absence of direct sequence conservation. By combining equivalent functional genomic data from two species, a synteny-based algorithm, and *in vivo* validation we reveal a substantial amount of previously hidden indirectly conserved elements functionally equivalent between mouse and chicken.

Identification of orthologous enhancers between distantly related species is an inherently difficult problem due to rapid enhancer evolution (Berthelot et al. 2017; Villar et al. 2015). While there have been several individual reports describing enhancers conserved in function rather than in sequence (Madgwick et al. 2019; Crocker and Stern 2017; Hare et al. 2008; Braasch et al. 2016; Fisher et al. 2006), a systematic evaluation of this phenomenon is challenging. Not only does it require algorithmic approaches that attempt to pair non-alignable sequences, but it also requires functional data that can be used to validate these predictions. By combining the synteny-based algorithm IPP with matching experimental data from two species, we were able to predict a large set of previously hidden indirectly conserved elements and demonstrate they are as likely to be functionally conserved as directly conserved elements. Our reanalysis of previous studies show that these likely 5-fold underestimate the number of chicken-conserved enhancers (Blow et al. 2010; Schmidt et al. 2010). While this does not change the general trend observed in these studies, the degree of underreported conserved regulatory elements changes the interpretation to which degree enhancers may evolve from neutral sequences (Galupa et al. 2023) and to which degree they are conserved. Our results indicate a degree of conservation invisible to current alignment-based measures. Thereby, our approach reconciles the apparent contrast between divergent non-coding genome sequences and other conserved features such as 3D chromatin structure and gene expression.

Rapidly diverging regulatory DNA allows adaptation of the regulatory genome during evolution but presents a major challenge for tracing the evolution of regulatory elements across species. Multiple sequence alignments and alignment-free algorithms are strategies to identify orthologous pairs of regulatory sequences, but are challenging, especially for large evolutionary distances. Efforts such as halliftover/HALPER (Zhang et al. 2020; Hickey et al. 2013) try to overcome this based on multiple alignment of hundreds of genomes. However, their performance is similar to IPP using only 16 genomes, highlighting the potential of synteny as a proxy for conservation. Bridged/tunneled alignments (Taher et al. 2011; Baranasic et al. 2022) provide a viable strategy for orthologous CRE detection and have already indicated a higher degree of CRE conservation than commonly assumed. Our approach builds on the idea of bridged alignments and extends it in several ways. One, IPP implements multiple bridging species, which can be optimized for any pairwise comparison based on their specific phylogenetic relationships. Two, within the framework of conserved synteny, IPP projections can assume orthology for any pair of regions between any two genomes, irrespective of their DNA sequence. Consequently, in non-syntenic regions, or between very distantly related genomes (Sanges et al. 2006) this strategy might miss orthologous elements. Nevertheless, IPP is a potent approach to identify putative orthologs for comparative studies at varying evolutionary distances provided the appropriate set of bridging species, in particular when combined with equivalent experimental data sets similar to our mouse and chicken heart data. Moreover, identification of indirectly conserved elements provides valuable information for interpretation of disease-associated non-coding variants in humans, for example in congenital heart disease (Richter et al. 2020; Xiao et al. 2024), and facilitates their functional characterization and testing in animal models.

Advances in machine learning have made it possible to predict the regulatory activity for any DNA sequence in a given cell type or tissue (Avsec et al. 2021; de Almeida et al. 2022; Reiter, de Almeida, and Stark 2023; de Almeida et al. 2023; Taskiran et al. 2023). Within mammals, models trained in one species can successfully predict activity in another (Minnoye et al. 2020; Kaplow et al. 2023; Kliesmete et al. 2024), but cannot match ortholog pairs. A recent study aimed to identify orthologous enhancers between human and mouse using a ML model, but requires syntenic regions as part of their algorithm to match orthologs (Oh and Beer 2023). Here we show that our SVM model trained in mouse can predict tissue-specific enhancers in chicken, highlighting the deep conservation of enhancer sequence syntax. Going beyond its predictive power, we use the model to independently validate IPP-projected regions in the chicken genome, demonstrating that indirectly conserved regions have sequence characteristics typical of heart enhancers. In the future, combination of both approaches can be a powerful strategy to study enhancer evolution. For example, IPP-identified pairs of orthologs can serve as training input for ML models to learn sequence changes compatible with functional conservation.

Sequence conservation of CREs, especially that of enhancers, displays a great level of heterogeneity ranging from ultra-conserved elements (Snetkova et al. 2021; Dickel et al. 2018; Snetkova et al. 2022) to the sequence-divergent IC elements we describe here. We show, however, that signals for functional conservation, in terms of chromatin signatures, encoded TFBS, and predicted tissue-specificity is relatively uncoupled from sequence conservation. As such, we imagine IPP to be an efficient approach to annotate orthologous CREs between species for example in single-cell ATAC-/ChIP-seq datasets from equivalent tissues, where cell types and expression programs are conserved, while the majority of CREs currently appear to be non-conserved.

Furthermore, the TFBS shuffling analysis suggests that CRE function may predominantly be maintained by TFBS composition. Consequently, conservation of an element’s regulatory function is less dependent on exact sequence conservation than on preserving the appropriate balance of TFBSs within the given element. Given that we found thousands of IC elements between mouse and chicken, the functional conservation of CREs across larger evolutionary distances is likely much more prevalent than currently appreciated.

## Code Availability

The source code for Interspecies Point Projection from this study can be obtained from https://github.com/tobiaszehnder/ipp

## Acknowledgements

D.M.I. and M.P. were supported by funding from the DFG SPP 22.02 “3D Genome Archictecture in Development and Disease” (IB139/1-1 and IB 139/6-1). Work in the Ibrahim Lab is supported by an ERC Starting Grant SYNREG (101076709). We thank the MPI-MG transgene facility and animal house for husbandry and members of Dominik Seelow and Martin Kircher’s labs for feedback on the Machine Learning analysis. We would like to thank Juliane Glaser, Alicia Madgwick and all members of the Ibrahim lab for feedback on the manuscript.

---

**Figure S1:**
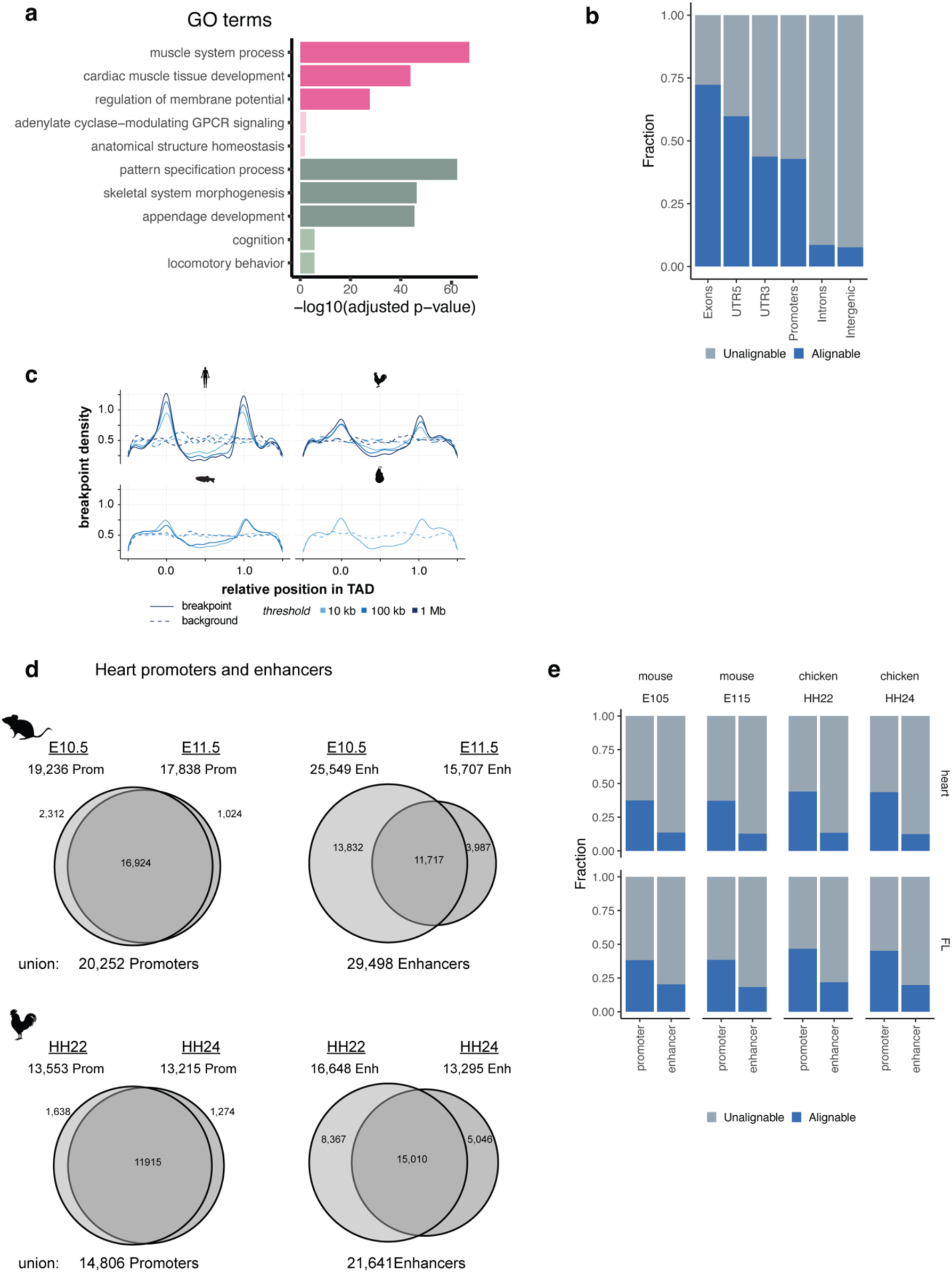
**(a)** Gene Ontology (GO) annotations of differentially expressed genes (Heart vs. FL) in mouse and chicken. Dark pink = upregulated, both species. Dark green = downregulated, both species. Light pink = upregulated, mouse-only. Light green = upregulated, chicken-only. Grey = no differential expression. **(b)** Estimation of sequence alignability of ATAC-seq peaks from mouse embryonic heart at different annotated genomic locations. **(c)** Number of predicted promoters and enhancers from stage-specific and shared/union sets in both species. **(d)** Estimation of sequence alignability from stage-specific predicted promoters and enhancers from heart and FL in both species.

**Figure S2:**
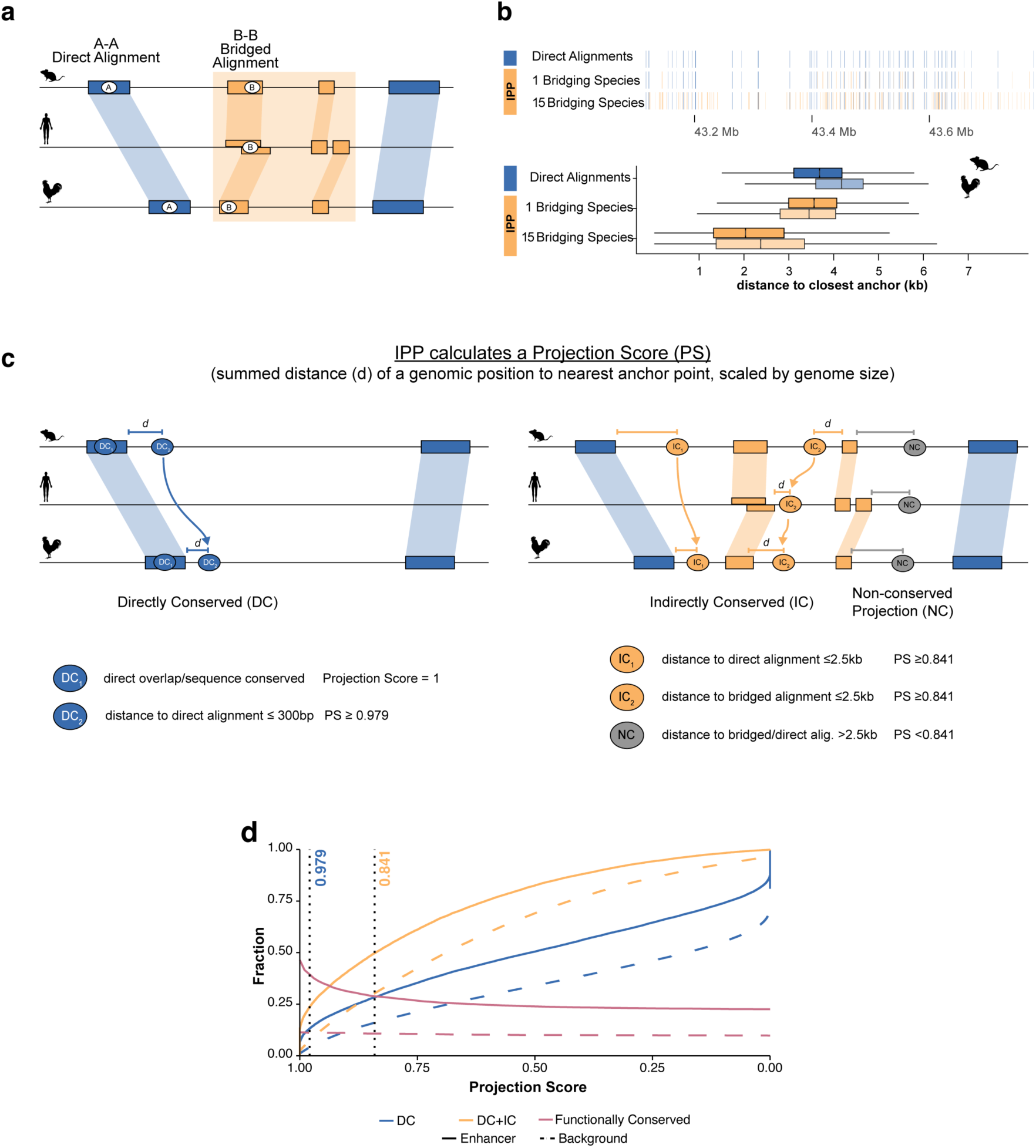
Interspecies Point Projection combines bridged alignments and synteny to identify orthologous regions. **(a)** Classification of direct and bridged alignments through the use of intermediate species **(b)** Increase in the number of anchor points and distance to the nearest anchor points through multi-species bridged alignments. Comparison between 0, 1 and 15 bridging species **(c)** Classifcation of projections as directly and indirectly conserved. DC regions overlap a sequence alignment or are ^≤^ 300bp from a direct alignment. The distance of IC regions as >300bp but ^≤^ 2.5kb from a direct or indirect alignment. Regions with >2.5kb summed distance through the species graph from anchor points are classified as NC. **(d)** Fractions of mouse enhancers identified as directly conserved (DC, blue) or either directly or indirectly conserved (DC + IC, orange) as a function of the projection score threshold. Fraction of functionally conserved DC+IC elements as a function of the projection score threshold (red). Solid lines = enhancers, dashed lines = randomly selected background regions. Dotted vertical lines represent DC threshold score of 0.979 and IC of 0.841.

**Figure S3.**
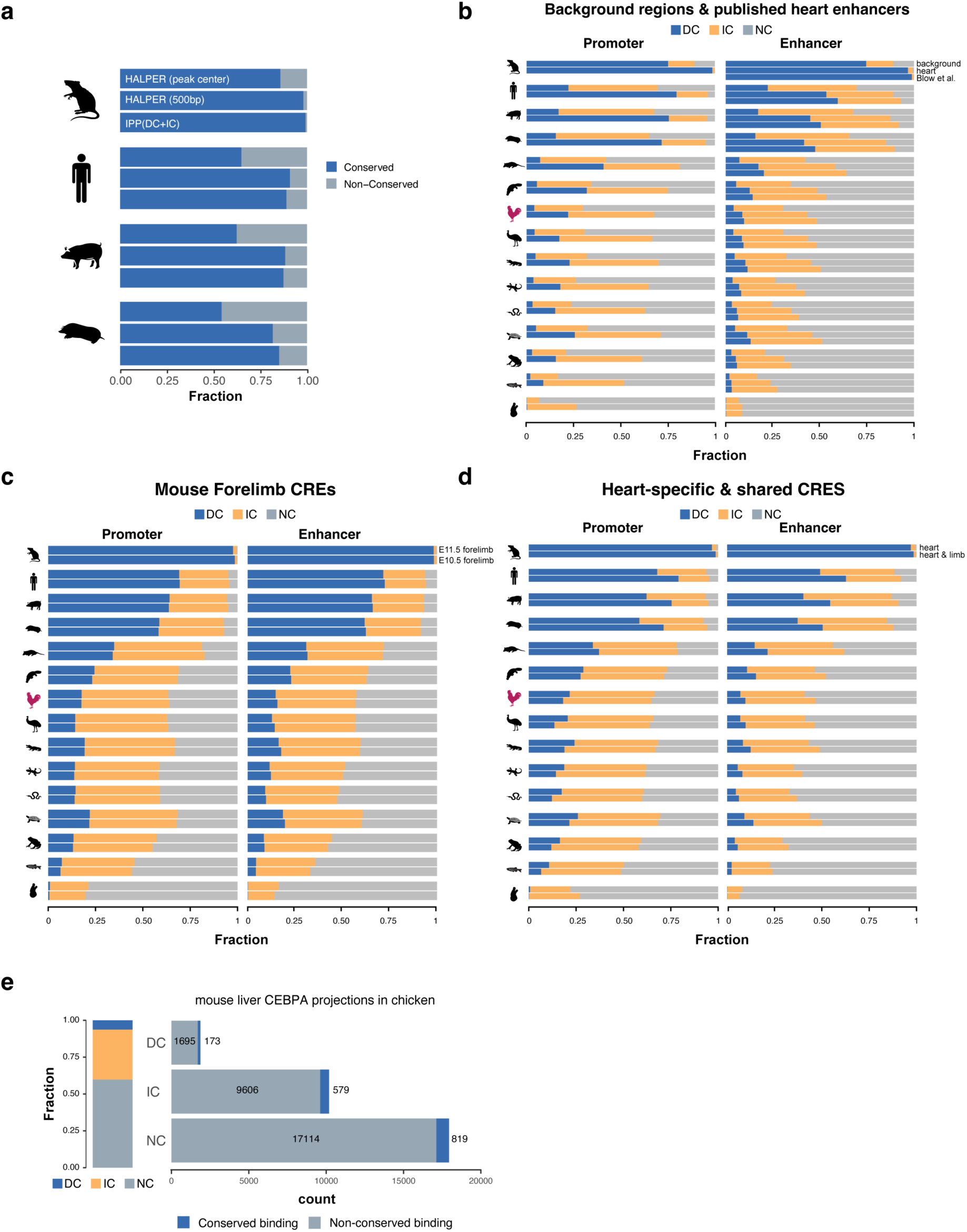
**(a)** IPP performance compared to halliftover/HALPER for mouse heart enhancer ortholog prediction in four placental mammals. **(b-d)** IPP projections for randomly selected genomic regions and published heart enhancers (Blow et al 2010) **(b)**, forelimb CREs at E10.5 & E11.5 **(c)**, and heart-specific or heart and limb CREs **(d)**. **(e)** Fraction of directly/indirectly conserved mouse CEBP/A ChIP-seq peaks in the chicken genome. Blue fractions (right) show the number of conserved binding events (as determined by overlap with a CEBP/A ChIP-seq peak in chicken livers)

**Figure S4.**
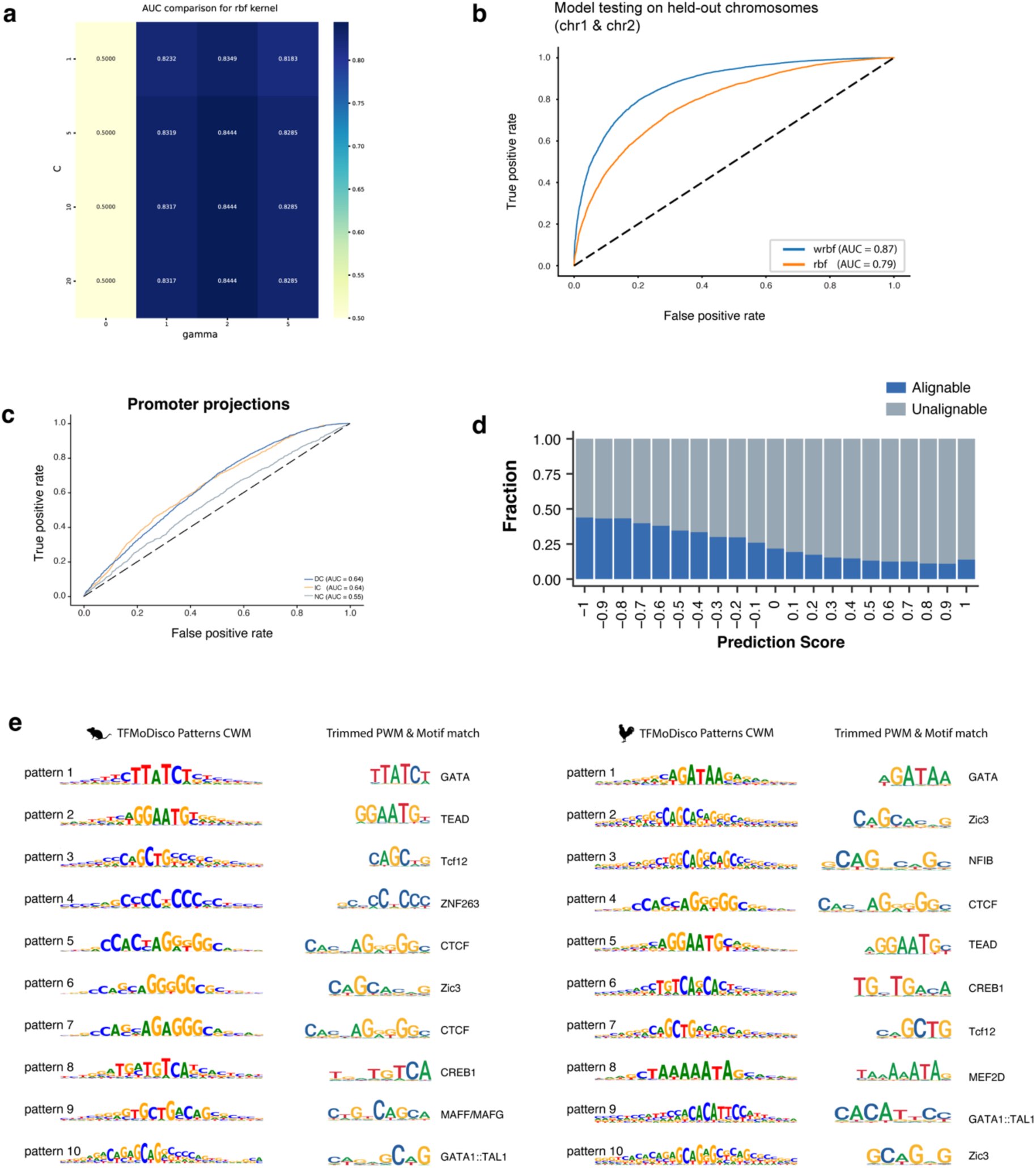
**(a)** Parameter tuning to train the SVM with RBF kernel with a grid-search for parameters c and gamma showing the calculated AUC after 5-fold cross validation. AUC = Area under the ROC curve. **(b)** ROC curves with computed AUC showing the performance of gkm-SVM with either RBF(rbf, orange) or weighted RBF(wrbf, blue) kernel on test data. The SVM was trained with the c & gamma parameters chosen in (a). **(c)** ROC curves with computed AUC showing human-chicken interspecies prediction accuracy for different conservation classes of mouse promoters projected to chicken. **(d)** Estimation of sequence alignability as a function of SVM predicted tissue-specificity (as prediction score) for ATAC-Seq peaks from chicken embryonic heart. **(e)** Top 10 mouse (left) and chicken (right) patterns discovered by TF-MoDisco showing seqlet as CWM, trimmed and converted PWMs and their annotated JASPAR motif match.

**Figure S5.**
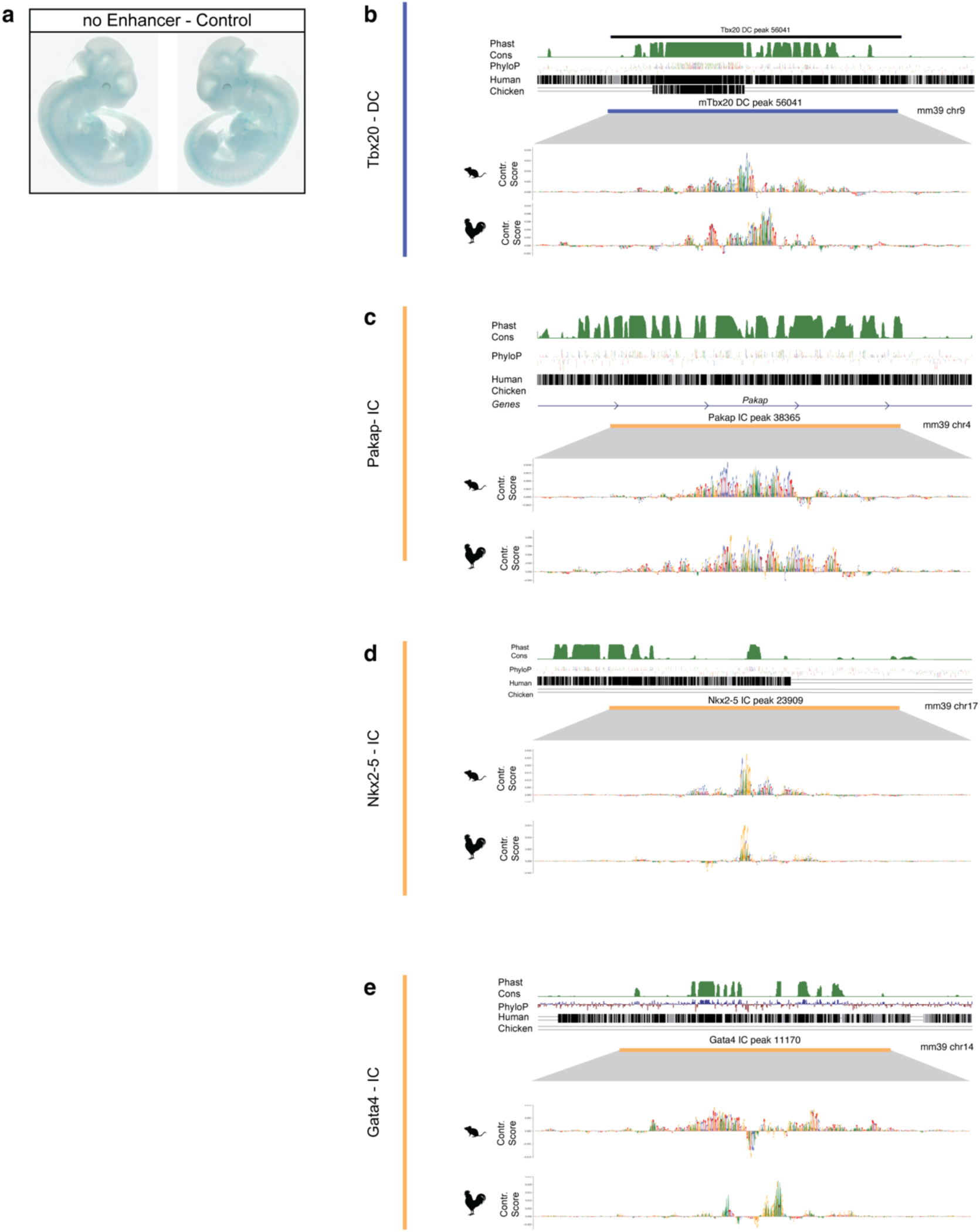
**(a)** Control for Enhancer reporter using a knock-in of the minimal promoter-lacZ without enhancer. Background signal in somites along the anterio-posterior axis. **(b-e)** Sequence conservation scores (PhastCons/PhyloP) and direct alignments to human and chicken of all tested enhancers. SVM contribution scores show important sequence features of the 500bp enhancers.

## Supplementary Text

### Interspecies Point Projection (IPP)

We project genomic point coordinates from a reference genome to a target genome by linear interpolation between blocks of pairwise sequence alignment, so called anchor points (Baranasic et al. 2022). Moreover, we use pairwise alignments between a set of bridging species to maximize anchor point density and thus optimize projection accuracy. This scenario is represented by a graph in which every node is a species, and the weighted edges represent the distance of a genomic coordinate to its anchor points between the nodes it connects (Fig. 2). We established a distance scoring function that returns a score of 1 for a genomic location *x* overlapping an anchor point (|*x* − *a*| = 0), and exponentially converges to zero with increasing distance |*x* − *a*|. For a single pairwise comparison, the function is defined as follows:

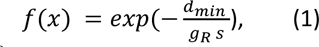

with 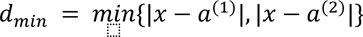 denoting the distance of a genomic location *x* to its closest anchor point, *g*_*R*_ denoting the genome size of the reference species and *s* a scaling factor that can be tweaked to determine the decreasing rate of the function. For instance, we can set s by defining a distance half life *d_h_* as the distance |*x* − *a*| at which the scoring function ought to return a value of 0.5:

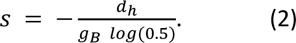

All projections presented in this manuscript were computed using a distance half life of 10 kb.

For the score calculation in Equation 1, the distance is normalized by the genome size of the reference species (*g*_*R*_) of a pairwise comparison. In Equation 2, the scaling factor is normalized by the size of a basis genome (*g*_*B*_) which we chose to be the mouse genome build mm39, allowing comparisons between projections from different reference species. In practice, this means that the distance scoring function decreases at equal rates for different reference genomes, however, these scores correspond to different distances based on the relative reference genome sizes. The function can thus be simplified to the following form:

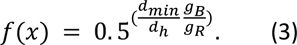

We can then compute the total distance score of a given path through the graph as the product of the score of all edges in that path. The length of a path is reciprocal to the distance scoring function, hence we can subtract the total score from 1 to obtain the path length *l*_*p*_:

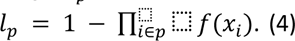

Finally, projection accuracy is optimized by finding the shortest path through the graph:

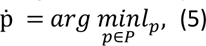

with *P* denoting the set of all paths through the graph connecting the reference and the target species. Finding the shortest path through a graph can be solved using Dijkstra’s Shortest Path Algorithm (Dijkstra 1959). We implemented the method in python and C++ and named it Interspecies Point Projection (IPP). IPP is publicly available at https://github.com/tobiaszehnder/ipp.

### Bridging species selection and pairwise alignment

IPP relies on the additional anchor points provided by bridging species to map corresponding genomic locations between pair of divergent genomes (**Fig. S2a,b**). As such, the choice of bridging species depends on the specific comparison of interest. Here, for a mouse-chicken comparison, we selected mammalian species which have diverged from chicken after mouse (human, pig, mole, opossum, platypus), and those which have diverged from mouse after chicken (alligator, green anole, snake, turtle) (**Supplementary Tab. 2**). Additionally, we selected the rat and emu as two closely related species to mouse and chicken, respectively. Finally, we included frogs, zebrafish, and the sea squirt as outgroups.

Fasta files for all reference genome assemblies were obtained either from DNA Zoo or from NBCI were used as inputs for pairwise alignments with *lastal*. Chain files were then generated, preprocessed, and merged for each species pair before combined in one collection of large pairwise alignments and stored in a binary format. This collection of alignments consisting of the reference, target, and all bridging species is the necessary input for running IPP.

### Projection classification and distance score tuning

IPP computes a score for every projection from one genome to another through the species graph. As described above, this score is a representation of the distance to the nearest anchor points, i.e. the higher the score, the shorter the distance and thus the more accurate the projection. We use this distance score as a threshold to classify projections into 3 classes: directly-conserved (DC), indirectly-conserved (IC), and non-conserved (NC) (**Fig. S2c**). An element is classified as *DC* if its projection score using only direct alignments was above this threshold, and as *IC* if their projection score from bridging alignment is also above such threshold. All remaining elements are then classified as *NC*. If ATAC-Seq data is available for the target genome, we further classify each projection by their functional conservation. Specifically, any projected point is classified as functionally conserved/’***+***’ or non-functionally conserved/’***-***‘, if it is within or outside a 2.5kb distance from an ATAC-seq peak summit, respectively.

Initially, we set the score threshold at 0.99 which, given Equation 2, represents a maximum distance to the next anchor point of ∼150 bps for DC elements. For IC elements, this additionally means that the sum of distances from the query element to an anchor point at all intermediate projections is =< 150bp. While ensuring the confidence of projections, this rather stringent cutoff implies a certain level of false negatives within NC, i.e. projections with a projection score below the cutoff that are nevertheless pointing to the correct ortholog. Furthermore, taking into practical consideration that IPP only maps a single base-pair of an element between genomes, such stringency likely results in an underestimation of conservation of the element of interest.

We then seek to tune this threshold parameter, which is ultimately a trade-off between specificity and sensitivity. In other words, relaxing the distance threshold will result in more elements being classified as conserved with a higher likelihood of such classification being a false positive (i.e. a projection pointing to a non-orthologous region. We took advantage of available ATAC-Seq data in the chicken forelimb as an independent tissue model, and determine if and how the proportion of functionally conserved elements changes as we relax the cut-off score. We observe a clear drop in the fraction of functionally conserved elements at high projection scores (i.e. >=0.9) from 38% to below 27% of all conserved enhancers (**Fig. S2d**). This sharp change in proportion appears to plateau at lower score thresholds. Indeed, even with dramatically forgiving cutoffs, just over 1/5th of all projections is putatively functionally conserved at every score threshold below 0.75. Importantly, this trend is not reflective of the spatial distribution of open chromatin, as only ∼10% of randomly selected background regions reside within open chromatin after projection (**Fig. S2d**).

One can take advantage of such a relaxed approach to identify putative *functionally* conserved orthologous enhancers (e.g. projected elements that are residing in open chromatin), providing an additional layer of functional validation. Given the availability of equivalent functional datasets, we decided to relax this cut-off and used 2 different distance cut-offs for DC and IC classifications. Specifically, an element with a projection score of 0.979 (∼300bp distance) using only direct alignments is classified as DC. For IC classification, we used a score cut-off of 0.841, which is equivalent to a summed distance of 2.5kb through all intermediate projections. These projections are filtered - as before - for those overlapping open chromatin regions to select for putatively functionally conserved elements. This permits the detection of functional orthologs in highly dynamic genomic neighborhoods where sequence alignments are sparse, with the potential cost of a higher false discovery rate.

### Materials and Methods Biological samples

C57/BL6 inbred mice were used for timed mating and fertilized SPF eggs (Valo Biomedia) were incubated at 38°C 50-55% humidity. Embryonic hearts and forelimbs from mouse and chicken embryos (E10.5, E11.5 and HH22, HH24) were dissected and further processed for sequencing libraries preparation. Each experiment was performed in biological replicates.

### Sample and Sequencing libraries preparation

#### a. RNA-seq

For RNA-seq, dissociated chicken embryonic heart cells were snap-frozen in liquid N2. RNA was extracted using the Qiagen RNeasy-Mini Kit according to manufacturer’s instructions. Ribosomal RNA was depleted before library preparation with the Kapa HyperPrep Kit and sequenced on a Novaseq2 100 bp paired-end reads. RNA-seq experiments were performed in duplicates.

#### b. ATAC-seq

ATAC-seq libraries were prepared using the Omni-ATAC protocol from 50k cells per replicate. Embryonic tissues were dissociated into single-cell suspension, washed with cold PBS, and lysed in fresh lysis buffer (10mM TrisCl pH7.4, 10mM NaCl, 3mM MgCl2, 0.1% (v/v) Igepal CA-630) on ice. Tn5 transposition for lysed nuclei was done for 30 min at 37° C, and DNA was then purified using the MinElute Reaction Cleanup kit (Qiagen) kit.

Nextera indexing primers were added during library amplification from purified DNA, where the number of cycles were determined by qPCR as described. After double-sided size selection, we verified the expected nucleosomal fragment distribution with a BioAnalyzer or TapeStation. DNA concentration of libraries were measured with Qubit HS before sequencing on a Novaseq2 (Illumina) using 100bp paired-end reads.

#### c. ChIPmentation

ChIPmentation libraries were prepared as previously described (Schmidl et al. 2015). Briefly, dissociated cells were first filtered through a 100µm (embryonic heart) or 70µm (limb) MACS® SmartStrainer before fixation with 1% MeOH-free formaldehyde (Thermo Scientific: 28906) in PBS on ice for 10 minutes. Fixed cells were first quenched using glycine and then lysed on ice in lysis buffer (10mM Tris pH 8.0, 100mM NaCl, 1mM EDTA pH 8.0, 0.5mM EGTA, 0.1% Sodium deoxycholate, 0.5% N-lauroylsarcosine) before shearing with a Covaris E220 for a fragment distribution of 200-700bp. Antibodies were incubated overnight at 4C, followed by immunoprecipitation with protein G beads (id). After beads washing, transposition/’tagmentation’ reaction with the Tn5 transposase was done at 37C for 5min. Beads were then again washed before overnight reverse crosslinking with Proteinase K. DNA was then purified using the MinElute Reaction Cleanup kit (Qiagen).

Libraries were indexed and amplified similarly as previously described for ATAC-Seq libraries. The number of PCR cycles for each library was estimated using Ct values as determined by qPCR (where number of cycles = rounded up Ct value +1). After amplification, DNA was cleaned up with AmPureXP beads, and then checked on a TapeStation D5000 HS for size distribution. Size selection was then carried out accordingly. The concentration of final eluted DNA was measured using Qubit HS and checked again on a TapeStation D5000HS. All libraries were sequenced on a Novaseq2 (Illumina) using 100bp paired-end reads.

#### d. Hi-C

In situ Hi-C libraries were prepared as previously described (Schöpflin et al. 2022). Briefly, 3C libraries were digested with DpnII, and digested ends were marked with biotin-14-dATP. DNA was sheared with an S-Series 220 Covaris to 300-600bp fragments before biotin pull-down using Dynabeads MyOne Streptavidin T1 beads. Sheared DNA ends were then repaired with T4 DNA polymerase and the Klenow fragment of DNA polymerase I, and subsequently and phosphorylated with T4 Polynucleotide Kinase NK. Sequencing adaptors were then added, and libraries were indexed via PCR amplification (4–8 cycles) using the NEBNext Ultra II Q5 Master Mix. PCR clean-up and size selection were done with AmPureXP beads before 100bp paired-end sequencing on a Novaseq2.

### Data processing

#### a. RNA

We processed all RNA-seq libraries with STARv2.7.9a using reference genome sequences and annotations from GENCODE (vM32, primary) for mouse and Ensembl (GRCg7b) for chicken. We obtained gene-level counts for each sample with *--quantMode geneCounts*. In addition to in-house chicken heart RNA-seq libraries, we similarly processed the following publicly available datasets: mouse heart E10.5 & E11.5 (ENCODE3), chicken FL HH22 & HH24 (<u>GSE164737,</u> (Jhanwar et al. 2021). TPM values were computed from gene-level counts, where gene length is estimated as the sum of all exon lengths.

#### b. ATAC-seq & ChIPmentation

For ATAC-seq & ChIPmentation samples, Nextera Tn5 adaptor sequences were trimmed from fastq reads using *cutadapt* before further processing. Reads were aligned to appropriate reference genomes (mm10, mm39, or galGal6) using bowtie2 *v2.3.5.1* where the maximum fragment length set was either 1000bp (ATAC) or 700bp (ChIPmentation). Duplicated reads were then removed using *MarkDuplicates* (Picard v2.23.4). Finally, reads were further sorted and filtered using samtools *v1.10* to remove unmapped reads, low quality reads (MAPQ < 10), and mitochondrial reads. Filtered bam files from replicates were merged to generate bigwig files. We used *bamCoverage* (deepTools) with CPM normalization and bin size of either 1 for ATAC or 10 for ChIPmentation.

Peak calling for ATAC-seq data from replicates was done with Genrich *v0.6.1* in ATAC mode ‘-j’ with default parameters. (https://github.com/jsh58/Genrich).

#### c. HiC

Reads handling were done using Juicer v1.6.0 CPU version (Durand NC, et al. 2016). Specifically, alignment was done with BWA-MEM v0.7.17 to reference genome galGal6. Only read pairs with MAPQ > 30 were included in the final contact maps. Processing was done separately for each replicate, and output filtered de-duplicated read pairs were merged. Contact matrices were balanced with Knight-Ruiz normalization (Knight PA & Ruiz D. 2012) before visualization.

### Data analysis

#### a. Comparative differential expression analysis

Raw gene-level counts from heart and limb samples at both stages were used as input for differential analysis with DESeq2 [v1.36](Love, Huber, and Anders 2014). We obtain a set of differentially expressed genes in the heart relative to limb in both stages, accounting for the effects for biological replicates. To aid visualization and gene ranking for Gene Ontology (GO) analysis, effect size shrinkage was done for the coefficient modeling tissue-specificity (i.e. *tissue_heart_vs_limb*).

Gene orthology annotations were obtained from Ensembl databases GRCm39 for mouse and GRCg7b for chicken. Duplicated annotations were filtered to retain only those with the highest GOC score. Only one-to-one orthologous genes (OGs) were used for all comparative analysis.

Gene Ontology (GO) analysis was done using R package clusterProfiler [v4.4.4] (Wu et al. 2021). Over-representation GO analysis of OGs was done given a background gene set of all detectably expressed mouse genes (i.e. raw counts >= 10). For statistical testing, testing gene-set sizes were set from a minimum of 5 to a maximum of 100 genes to allow focusing of specific biological processes (i.e. BP) over more general terms.

#### b. Estimation of sequence alignability

To estimate conservation by means of sequence alignability, we used UCSC LiftOver as implemented within R package *rtracklayer* for reciprocal mapping between mouse and chicken genomes. Chain files for mm39 and galGal6 were obtained from UCSC before importing into R using *rtracklayer*. For mapping, we used the default parameter settings (minMatch=0.1) and allowed for multiple mapping (i.e. one-to-many) between query and target.

#### c. Enhancer & Promoter prediction

Histone profiles (i.e. H3K27ac, H3K4me1, H3K4me3) from merged replicates were used to predict candidate regulatory regions using the enhancer prediction tool CRUP (Ramisch et al. 2019). In brief, CRUP computes the probability score for each 100bp bin in the entire genome to be an active enhancer element. Combining these probabilities and normalized histone signal values (i.e. mono:tri ratios), bins are filtered and merged into either promoter-like or enhancer-like regions.

To define active promoter regions, we intersected defined promoter-like regions with all TSS of actively transcribed genes (i.e. counts ≥ 1TPM). Counts values were obtained as described previously for expression analysis. Once promoters are defined, we finalized the set of active enhancers by filtering enhancer-like regions by their accessibility from called ATAC peaks. Finally, those falling within 2kb of a predicted promoter are removed from the final set of active enhancers. The numbers of enhancers and promoters can be found in Sup. Tab. 1 and the bed files under GSE263587, GSE263753, GSE263755, GSE263783.

### TFBS Motif and Foot-printing Analysis

#### a. Reference motif collection

We obtained TF motif models from the JASPAR 2022 database (core vertebrate, non-redundant) and systematically curate this database to be used as a reference for all TFBS-based analysis. From over 700 JASPAR TF motifs, we filtered for those with detectable expression in the mouse embryonic heart by integrating RNA-seq counts (described above). Detectable expression is defined as having counts of >= 1 TPM in both replicates, in either stage E10.5 or E11.5 (n=520). From these, we further consolidate the reference collection by filtering out redundant motifs based on sequence similarity within the same annotated TF family. Specifically, within each TF family, motifs are ranked by their informational content score before pair-wise comparison with others in the same family using the *compare_motifs* function from R package *universalmotif*. Finally, motifs with lower informational content score and a similarity score > 0.9 (score of 1 = identical sequence) are discarded from the final reference set (n=301).

#### b. Motif scanning

To characterize the TFBS composition of CREs, we searched for motif matches from the curated collection using FIMO implemented through R package *memes* (Grant, Bailey, and Noble 2011) with default parameters. DNA sequences were obtained from annotation packages *BSgenome.Mmusculus.UCSC.mm39 BSgenome.Ggallus.UCSC.galGal6* for mouse and chicken, respectively. Motifs scanning was done within a 500bp window centered by ATAC peak summit or projected point. Peak centering by summit was done for projected regions in chicken only for functionally conserved elements (i.e. DC+ & IC+). Finally, any overlapping hits from the same motifs are discarded, keeping the match with higher score.

#### c. ATAC-seq foot-printing

Aligned ATAC-seq reads from biological replicates were merged to be used as input for ATAC-seq footprinting analysis using TOBIAS (Bentsen et al. 2020) **[v0.3.3]**. The genomic regions of interests to be foot-printed were: (1) the union set of predicted enhancers and promoters in mouse and chicken hearts, and (2) all called chicken ATAC-seq peaks. Briefly, we used TOBIAS to correct for Tn5 bias before footprint scores were calculated at genomic regions of interest. Finally, we used our curated set of TFSB motifs as reference to predict TF binding. TOBIAS output from different stages were merged, and overlapping regions of predicted binding from the same TF were merged similarly to motif hits as described. Finally, quantification of shared footprints was done similarly to the motifs analysis previously described.

#### d. Quantification of motifs and TFBS sharing between pairs of orthologous CREs

To quantify the similarity between mouse CREs and their corresponding chicken orthologs as determined by IPP, we determine the total number of shared motifs and TF-binding (i.e. TFBS) between every mouse-chicken pair of sequences. As a negative control, we also compare the number of shared motifs and TFBS between a mouse sequence and non-orthologous, i.e. background genomic region. Specifically, for every mouse sequence with a chicken projection overlapping an ATAC-seq peak (i.e DC+/IC+/NC+), another ATAC-seq peak (if possible, within the same TAD) is randomly selected as its non-ortholog.

### Classification model for heart-specific enhancers

#### a. Training strategy and data preparation

Our classification model is a Support Vector Machine (SVM) with a center-weighted radial basis gapped k-mer kernel function (wrbfgkm) (implemented at https://github.com/kundajelab/lsgkm-svr) (Ghandi et al. 2014; Lee 2016). All datasets used for model training are processed bulk ATAC-seq data either obtained from ENCODE or in-house (as described above). To learn predictive features of heart-specific enhancers, we construct the positive set to include called ATAC-seq peaks from mouse hearts at 6 developmental stages (**in-house**: E10.5 & E11.5, **ENCODE**: E12.5-E14.5 & P0). All regions are centered at peak summit and extending 250bp on either side. Additionally, to ensure the model learns enhancer-specific regulatory features, regions within 2kb of an annotated mouse promoters (from EPD3 database) were removed from the final training set (n=∼65k).

For model training, we construct the negative set such that the model can accurately learn the sequence features determining whether an enhancer/CRE is heart-specific. First, to limit confounding factors, we generated a 10-fold null set of from random genomic loci. From these regions, we filtered for those overlapping any annotated ENCODE candidate CREs or ATAC-seq peaks from 5 non-heart embryonic organs (limbs, mid-/fore/hind-brain, liver, E12.5) and mESCs. Finally, those within a 2-kb overlap of any regions from the positive set were removed (n=70k).

All negative sets of GC- and repeats-matched sequences were generated using the *genNullSeqs* function from R package gkmSVM (Ghandi et al. 2014, 2016). Repeats-masked genomic sequences were obtained from custom masked *BSgenome* data packages for mm10, mm39 or galGal6.

#### b. Hyperparameter tuning & performance evaluation

As a measure for classification performance, the area under the ROC curve (AUC) was computed and visualized. For parameter tuning, a grid search for *C* and *g* parameters for wrbfgkm-kernel was done using a 5-fold cross validation for each combination of *C*= 1, 5, 10, 20 and *g* = 0, 1, 2, 5 (=16 conditions). The best performing parameter set (c=10, g=2) as determined by its calculated AUC was chosen for model training. The final model was tested on positive vs. negative regions on held-out chromosome 1 & 2.

#### c. Model prediction on chicken CREs and projections

Our heart-enhancer SVM model trained on mouse sequences was used to classify: (1) identified chicken enhancer and promoter sequences (described previously) from heart and FL, and (2) sequences mouse CREs at the projected chicken regions from by IPP. For each prediction, the negative set generated as described previously consists of GC- and repeats-matched regions. Additionally, only projected regions overlapping an ATAC-seq peaks (i.e. DC+ or IC+) were included in the analysis. AUROCs were computed to evaluate the model’s performance on these regions.

#### d. Model interpretation and de novo motifs discovery

We used GkmExplain (Shrikumar, Prakash, and Kundaje 2019) (implemented at https://github.com/kundajelab/lsgkm-svr) to interpret the model’s classification. GkmExplain computes the contribution score at each nucleotide to the SVM classification in all input sequences, i.e. its *importance score*. For each sequence, this importance score was computed by element-wise multiplication of the one-hot encoded sequence matrix by its hypothetical importance score. Scores were visualized using the *visualization* module from Python package *modisco*.

Computed hypothetical score was then normalized by the ratio of original importance scores and sum of all hypothetical scores having the same sign. Normalization allows the score to better reflect the importance of a specific base at each position thereby reducing noise for subsequent motif discovery with TF-Modisco (Shrikumar et al. 2018) (implemented at https://github.com/jmschrei/tfmodisco-lite). Computed and normalized scores from GkmExplain from: (1) mouse positive test set (n=9k), and (2) heart-specific chicken enhancers (n=15k) were used as input for two separate TF-Modisco runs. Similar positive sequence patterns from these were then merged for the final set of predictive sequence patterns and stored as PWM motifs. Flanking positions with information content < 0.5 were trimmed from the PWMs before being annotated with known motifs using TOMTOM (Gupta et al. 2007) with our TF motifs collection as reference.

### Quantification of motifs shuffling

To quantify the degree of motifs shuffling, we measure the Kendall tau distance (K_d_) between pairs of reference mouse sequence and its corresponding chicken orthologs. The Kendall tau distance metric measures the similarity between two ranking lists by counting the number of transpositions, or swaps, needed between pair of ranks in one list to achieve the same order from another list. The more similar the two lists are, the smaller the distance. A pair of mouse-chicken sequences is considered two ranking lists of motifs, where the order of shared motifs is the ranks. Here, we encode only the 5’-3’ order of motifs for mouse sequences as reference and compare them to both orientations of the chicken sequences.

To ensure we faithfully encode the specific order of motifs as ranks, shared motifs obtained previously are further processed to filter out largely overlapping occurrences from different motifs (minimum overlap of 8bp), again keeping the hits with the highest mapping score. Additionally, to ensure unique rankings, runs of hits from the same motif are considered a singular match. Any sequence containing >1 noncontiguous hits from the same motif (e.g. A,B,C,A,D) is stored as a matrix of ranking lists, where each row represents a unique ranking order (e.g. 1-A,B,C,D and 2-B,C,A,D). Using R package *rankdist* (Qian and Yu 2019), we compute the normalized K_D_ between all unique ranking lists for a mouse-chicken pair, which accounts for varying number of shared motifs (i.e. list length). Finally, assuming the fewest possible changes have occurred during evolution, we take the smallest computed K_d_ value for every pairwise comparison and compared between conservation classes DC, IC, and NC. We also described the effect size of sequence conservation on sequence shuffling by computing Cohen’s d using R package *effsize* (Torchiano 2016).

### In vivo enhancer-reporter assays

Transgenic mice carrying the individual mouse or chicken enhancers tested in this study were generated using a site-specific integration protocol, modified for mouse embryonic stem cells (mESCs). The PhiC31 system used (Chi et al. 2019) allows precise recombination between two att sites: the attP site inserted in a safe harbour genomic locus and the attB site in a donor vector. Genomic regions and primers used for generation of Enhancer Reporters can be found in Supplementary Table 2.

First, a master mESC line was established in which an Hsp68::LacZ expression cassette containing the attP site was inserted into a safe harbour locus (H11) via CRISPR/Cas9 using FuGENE technology (Promega). To create the donor vectors, we cloned each individual enhancer in a vector containing the attB site and a puromycin (Sigma-Aldrich, P8833) selection marker using Gibson cloning. Subsequently, each resulting donor vector was co-transfected with the PhiC31 plasmid into the master line using Lipofectamine LTX (Invitrogen), following the manufacturer’s guidelines. The enhancer-reporter mESC lines were cultured and embryos were generated via tetraploid complementation (Artus and Hadjantonakis 2011). At embryonic day E10.5, the embryos were harvested and processed for LacZ staining. Briefly, the embryos were kept in the dark at 37°C in LacZ staining buffer supplemented with 0.5 mg/ml X-gal, 5 mM potassium ferrocyanide and 5 mM potassium ferricyanide. When the desired staining was achieved, the embryos were washed several times in PBS and then fixed with 4% PFA/PBS supplemented with 0.2% glutaraldehyde and 5mM EDTA for long-term storage at 4°C. Embryos were imaged using a SteREO Discovery.V12 microscope with CL9000 cold light source and a Leica DFC420 digital camera. The embryo genotyping was performed by PCR using primers spanning the expected 5’ and 3’ integration junctions to confirm correct integration of the enhancers.

## References

Almeida, Bernardo P. de, Franziska Reiter, Michaela Pagani, and Alexander Stark. 2022. “DeepSTARR Predicts Enhancer Activity from DNA Sequence and Enables the de Novo Design of Synthetic Enhancers.” Nature Genetics 54 (5): 613–24.

Almeida, Bernardo P. de, Christoph Schaub, Michaela Pagani, Stefano Secchia, Eileen E. M. Furlong, and Alexander Stark. 2023. “Targeted Design of Synthetic Enhancers for Selected Tissues in the Drosophila Embryo.” Nature, December, 1–5.

Armstrong, Joel, Glenn Hickey, Mark Diekhans, Ian T. Fiddes, Adam M. Novak, Alden Deran, Qi Fang, et al. 2020. “Progressive Cactus Is a Multiple-Genome Aligner for the Thousand-Genome Era.” Nature 587 (7833): 246–51.

Artus, Jérôme, and Anna-Katerina Hadjantonakis. 2011. “Generation of Chimeras by Aggregation of Embryonic Stem Cells with Diploid or Tetraploid Mouse Embryos.” In Transgenic Mouse Methods and Protocols, edited by Marten H. Hofker and Jan van Deursen, 693:37–56. Methods in Molecular Biology. Humana Press.

Avsec, Žiga, Vikram Agarwal, Daniel Visentin, Joseph R. Ledsam, Agnieszka Grabska-Barwinska, Kyle R. Taylor, Yannis Assael, John Jumper, Pushmeet Kohli, and David R. Kelley. 2021. “Effective Gene Expression Prediction from Sequence by Integrating Long-Range Interactions.” Nature Methods 18 (10): 1196–1203.

Baranasic, Damir, Matthias Hörtenhuber, Piotr J. Balwierz, Tobias Zehnder, Abdul Kadir Mukarram, Chirag Nepal, Csilla Várnai, et al. 2022. “Multiomic Atlas with Functional Stratification and Developmental Dynamics of Zebrafish Cis-Regulatory Elements.” Nature Genetics 54 (7): 1037–50.

Bejerano, G., A. C. Siepel, W. J. Kent, and D. Haussler. 2005. “Computational Screening of Conserved Genomic DNA in Search of Functional Noncoding Elements.” Nature Methods 2 (7): 535–45.

Bentsen, Mette, Philipp Goymann, Hendrik Schultheis, Kathrin Klee, Anastasiia Petrova, René Wiegandt, Annika Fust, et al. 2020. “ATAC-Seq Footprinting Unravels Kinetics of Transcription Factor Binding during Zygotic Genome Activation.” Nature Communications 11 (1): 4267.

Berthelot, Camille, Diego Villar, Julie E. Horvath, Duncan T. Odom, and Paul Flicek. 2017. “Complexity and Conservation of Regulatory Landscapes Underlie Evolutionary Resilience of Mammalian Gene Expression.” Nature Ecology & Evolution 2 (1): 152–63.

Blow, Matthew J., David J. McCulley, Zirong Li, Tao Zhang, Jennifer A. Akiyama, Amy Holt, Ingrid Plajzer-Frick, et al. 2010. “ChIP-Seq Identification of Weakly Conserved Heart Enhancers.” Nature Genetics 42 (9): 818–22.

Braasch, Ingo, Andrew R. Gehrke, Jeramiah J. Smith, Kazuhiko Kawasaki, Tereza Manousaki, Jeremy Pasquier, Angel Amores, et al. 2016. “The Spotted Gar Genome Illuminates Vertebrate Evolution and Facilitates Human-Teleost Comparisons.” Nature Genetics 48 (4): 427–37.

Chi, Xiuling, Qi Zheng, Ruhong Jiang, Ruby Yanru Chen-Tsai, and Ling-Jie Kong. 2019. “A System for Site-Specific Integration of Transgenes in Mammalian Cells.” PloS One 14 (7): e0219842.

Crocker, J., and D. L. Stern. 2017. “Functional Regulatory Evolution Outside of the Minimal Even-Skipped Stripe 2 Enhancer.” Development 144 (17): 3095–3101.

Dickel, Diane E., Athena R. Ypsilanti, Ramón Pla, Yiwen Zhu, Iros Barozzi, Brandon J. Mannion, Yupar S. Khin, et al. 2018. “Ultraconserved Enhancers Are Required for Normal Development.” Cell 172 (3): 491–499.e15.

Dijkstra, E. W. 1959. “A Note on Two Problems in Connexion with Graphs.” Numerische Mathematik 1 (1): 269–71.

Engström, Pär G., Shannan J. Ho Sui, Oyvind Drivenes, Thomas S. Becker, and Boris Lenhard. 2007. “Genomic Regulatory Blocks Underlie Extensive Microsynteny Conservation in Insects.” Genome Research 17 (12): 1898–1908.

Firulli, A. B., D. G. McFadden, Q. Lin, D. Srivastava, and E. N. Olson. 1998. “Heart and Extra-Embryonic Mesodermal Defects in Mouse Embryos Lacking the BHLH Transcription Factor Hand1.” Nature Genetics 18 (3): 266–70.

Fisher, Shannon, Elizabeth A. Grice, Ryan M. Vinton, Seneca L. Bessling, and Andrew S. McCallion. 2006. “Conservation of RET Regulatory Function from Human to Zebrafish without Sequence Similarity.” Science (New York, N.Y.) 312 (5771): 276–79.

Galupa, Rafael, Gilberto Alvarez-Canales, Noa Ottilie Borst, Timothy Fuqua, Lautaro Gandara, Natalia Misunou, Kerstin Richter, et al. 2023. “Enhancer Architecture and Chromatin Accessibility Constrain Phenotypic Space during Drosophila Development.” Developmental Cell 58 (1): 51–62.e4.

Ghandi, Mahmoud, Dongwon Lee, Morteza Mohammad-Noori, and Michael A. Beer. 2014. “Enhanced Regulatory Sequence Prediction Using Gapped K-Mer Features.” PLoS Computational Biology 10 (7): e1003711.

Ghandi, Mahmoud, Morteza Mohammad-Noori, Narges Ghareghani, Dongwon Lee, Levi Garraway, and Michael A. Beer. 2016. “GkmSVM: An R Package for Gapped-Kmer SVM.” Bioinformatics (Oxford, England) 32 (14): 2205–7.

Grant, Charles E., Timothy L. Bailey, and William Stafford Noble. 2011. “FIMO: Scanning for Occurrences of a given Motif.” Bioinformatics 27 (7): 1017–18.

Gupta, Shobhit, John A. Stamatoyannopoulos, Timothy L. Bailey, and William Stafford Noble. 2007. “Quantifying Similarity between Motifs.” Genome Biology 8 (2): R24.

Hare, Emily E., Brant K. Peterson, Venky N. Iyer, Rudolf Meier, and Michael B. Eisen. 2008. “Sepsid Even-Skipped Enhancers Are Functionally Conserved in Drosophila despite Lack of Sequence Conservation.” PLoS Genetics 4 (6): e1000106.

Harmston, Nathan, Elizabeth Ing-Simmons, Ge Tan, Malcolm Perry, Matthias Merkenschlager, and Boris Lenhard. 2017. “Topologically Associating Domains Are Ancient Features That Coincide with Metazoan Clusters of Extreme Noncoding Conservation.” Nature Communications 8 (1): 441.

Heikinheimo, M., J. M. Scandrett, and D. B. Wilson. 1994. “Localization of Transcription Factor GATA-4 to Regions of the Mouse Embryo Involved in Cardiac Development.” Developmental Biology 164 (2): 361–73.

Hickey, Glenn, Benedict Paten, Dent Earl, Daniel Zerbino, and David Haussler. 2013. “HAL: A Hierarchical Format for Storing and Analyzing Multiple Genome Alignments.” Bioinformatics (Oxford, England) 29 (10): 1341–42.

Irie, Naoki, and Shigeru Kuratani. 2011. “Comparative Transcriptome Analysis Reveals Vertebrate Phylotypic Period during Organogenesis.” Nature Communications 2 (1): 248.

Jhanwar, Shalu, Jonas Malkmus, Jens Stolte, Olga Romashkina, Aimée Zuniga, and Rolf Zeller. 2021. “Conserved and Species-Specific Chromatin Remodeling and Regulatory Dynamics during Mouse and Chicken Limb Bud Development.” Nature Communications 12 (1): 5685.

Jin, Sheng Chih, Jason Homsy, Samir Zaidi, Qiongshi Lu, Sarah Morton, Steven R. DePalma, Xue Zeng, et al. 2017. “Contribution of Rare Inherited and de Novo Variants in 2,871 Congenital Heart Disease Probands.” Nature Genetics 49 (11): 1593–1601.

Kaplow, Irene M., Alyssa J. Lawler, Daniel E. Schäffer, Chaitanya Srinivasan, Heather H. Sestili, Morgan E. Wirthlin, Badoi N. Phan, et al. 2023. “Relating Enhancer Genetic Variation across Mammals to Complex Phenotypes Using Machine Learning.” Science (New York, N.Y.) 380 (6643): eabm7993.

Kikuta, H., M. Laplante, P. Navratilova, A. Z. Komisarczuk, P. G. Engström, D. Fredman, A. Akalin, et al. 2007. “Genomic Regulatory Blocks Encompass Multiple Neighboring Genes and Maintain Conserved Synteny in Vertebrates.” Genome Research 17 (5): 545–55.

Kliesmete, Zane, Peter Orchard, Victor Yan Kin Lee, Johanna Geuder, Simon M. Krauß, Mari Ohnuki, Jessica Jocher, Beate Vieth, Wolfgang Enard, and Ines Hellmann. 2024. “Evidence for Compensatory Evolution within Pleiotropic Regulatory Elements.” BioRxiv. 10.1101/2024.01.10.575014.

Kuhn, R. M., D. Karolchik, A. S. Zweig, T. Wang, K. E. Smith, K. R. Rosenbloom, B. Rhead, et al. 2009. “The UCSC Genome Browser Database: Update 2009.” Nucleic Acids Research 37 (Database issue): D755-61.

Lee, Dongwon. 2016. “LS-GKM: A New Gkm-SVM for Large-Scale Datasets.” Bioinformatics (Oxford, England) 32 (14): 2196–98.

Love, Michael, Wolfgang Huber, and Simon Anders. 2014. “Moderated Estimation of Fold Change and Dispersion for RNA-Seq Data with DESeq2.” Genome Biology 15 (12): 550.

Ludwig, M. Z., N. H. Patel, and M. Kreitman. 1998. “Functional Analysis of Eve Stripe 2 Enhancer Evolution in Drosophila: Rules Governing Conservation and Change.” Development (Cambridge, England) 125 (5): 949–58.

Madgwick, Alicia, Marta Silvia Magri, Christelle Dantec, Damien Gailly, Ulla-Maj Fiuza, Léo Guignard, Sabrina Hettinger, Jose Luis Gomez-Skarmeta, and Patrick Lemaire. 2019. “Evolution of Embryonic Cis-Regulatory Landscapes between Divergent Phallusia and Ciona Ascidians.” Developmental Biology 448 (2): 71–87.

Maric, Darko, Aleksandra Paterek, Marion Delaunay, Irene Pérez López, Miroslav Arambasic, and Dario Diviani. 2021. “A-Kinase Anchoring Protein 2 Promotes Protection against Myocardial Infarction.” Cells (Basel, Switzerland) 10 (11): 2861.

McFadden, D. G., J. Charité, J. A. Richardson, D. Srivastava, A. B. Firulli, and E. N. Olson. 2000. “A GATA-Dependent Right Ventricular Enhancer Controls DHAND Transcription in the Developing Heart.” Development (Cambridge, England) 127 (24): 5331–41.

McGaughey, David M., Ryan M. Vinton, Jimmy Huynh, Amr Al-Saif, Michael A. Beer, and Andrew S. McCallion. 2008. “Metrics of Sequence Constraint Overlook Regulatory Sequences in an Exhaustive Analysis at Phox2b.” Genome Research 18 (2): 252–60.

Minnoye, Liesbeth, Ibrahim Ihsan Taskiran, David Mauduit, Maurizio Fazio, Linde Van Aerschot, Gert Hulselmans, Valerie Christiaens, et al. 2020. “Cross-Species Analysis of Enhancer Logic Using Deep Learning.” Genome Research 30 (12): 1815–34.

Oh, Jin Woo, and Michael A. Beer. 2023. “Gapped-Kmer Sequence Modeling Robustly Identifies Regulatory Vocabularies and Distal Enhancers Conserved between Evolutionarily Distant Mammals.” BioRxiv. 10.1101/2023.10.06.561128.

Olson, E. N. 2006. “Gene Regulatory Networks in the Evolution and Development of the Heart.”

Overbeek, P. A. 1997. “Right and Left Go DHAND and EHAND.” Nature Genetics. Springer Science and Business Media LLC.

Paten, Benedict, Dent Earl, Ngan Nguyen, Mark Diekhans, Daniel Zerbino, and David Haussler. 2011. “Cactus: Algorithms for Genome Multiple Sequence Alignment.” Genome Research 21 (9): 1512–28.

Pediatric Cardiac Genomics Consortium, Bruce Gelb, Martina Brueckner, Wendy Chung, Elizabeth Goldmuntz, Jonathan Kaltman, Juan Pablo Kaski, et al. 2013. “The Congenital Heart Disease Genetic Network Study: Rationale, Design, and Early Results.” Circulation Research 112 (4): 698–706.

Prall, Owen W. J., Mary K. Menon, Mark J. Solloway, Yusuke Watanabe, Stéphane Zaffran, Fanny Bajolle, Christine Biben, et al. 2007. “An Nkx2-5/Bmp2/Smad1 Negative Feedback Loop Controls Heart Progenitor Specification and Proliferation.” Cell 128 (5): 947–59.

Qian, Zhaozhi, and Philip L. H. Yu. 2019. “Weighted Distance-Based Models for Ranking Data Using the R Package Rankdist.” Journal of Statistical Software 90 (5). 10.18637/jss.v090.i05.

Ramisch, Anna, Verena Heinrich, Laura V. Glaser, Alisa Fuchs, Xinyi Yang, Philipp Benner, Robert Schöpflin, et al. 2019. “CRUP: A Comprehensive Framework to Predict Condition-Specific Regulatory Units.” Genome Biology 20 (1): 227.

Reiter, Franziska, Bernardo P. de Almeida, and Alexander Stark. 2023. “Enhancers Display Constrained Sequence Flexibility and Context-Specific Modulation of Motif Function.” Genome Research 33 (3): 346–58.

Richter, Felix, Sarah U. Morton, Seong Won Kim, Alexander Kitaygorodsky, Lauren K. Wasson, Kathleen M. Chen, Jian Zhou, et al. 2020. “Genomic Analyses Implicate Noncoding de Novo Variants in Congenital Heart Disease.” Nature Genetics 52 (8): 769–77.

Sanges, Remo, Eva Kalmar, Pamela Claudiani, Maria D’Amato, Ferenc Muller, and Elia Stupka. 2006. “Shuffling of Cis-Regulatory Elements Is a Pervasive Feature of the Vertebrate Lineage.” Genome Biology 7 (7): R56.

Schmidl, Christian, André F. Rendeiro, Nathan C. Sheffield, and Christoph Bock. 2015. “ChIPmentation: Fast, Robust, Low-Input ChIP-Seq for Histones and Transcription Factors.” Nature Methods 12 (10): 963–65.

Schmidt, D., M. D. Wilson, B. Ballester, P. C. Schwalie, G. D. Brown, A. Marshall, C. Kutter, et al. 2010. “Five-Vertebrate ChIP-Seq Reveals the Evolutionary Dynamics of Transcription Factor Binding.” Science 328 (5981): 1036–40.

Schöpflin, Robert, Uirá Souto Melo, Hossein Moeinzadeh, David Heller, Verena Laupert, Jakob Hertzberg, Manuel Holtgrewe, et al. 2022. “Integration of Hi-C with Short and Long-Read Genome Sequencing Reveals the Structure of Germline Rearranged Genomes.” Nature Communications 13 (1): 6470.

Shrikumar, Avanti, Eva Prakash, and Anshul Kundaje. 2019. “GkmExplain: Fast and Accurate Interpretation of Nonlinear Gapped k-Mer SVMs.” Bioinformatics (Oxford, England) 35 (14): i173–82.

Shrikumar, Avanti, Katherine Tian, Žiga Avsec, Anna Shcherbina, Abhimanyu Banerjee, Mahfuza Sharmin, Surag Nair, and Anshul Kundaje. 2018. “Technical Note on Transcription Factor Motif Discovery from Importance Scores (TF-MoDISco) Version 0.5.6.5.” ArXiv [Cs.LG]. arXiv. http://arxiv.org/abs/1811.00416.

Siepel, A., G. Bejerano, J. S. Pedersen, A. S. Hinrichs, M. Hou, K. Rosenbloom, H. Clawson, et al. 2005. “Evolutionarily Conserved Elements in Vertebrate, Insect, Worm, and Yeast Genomes.” Genome Research 15 (8): 1034–50.

Snetkova, Valentina, Len A. Pennacchio, Axel Visel, and Diane E. Dickel. 2022. “Perfect and Imperfect Views of Ultraconserved Sequences.” Nature Reviews. Genetics 23 (3): 182–94.

Snetkova, Valentina, Athena R. Ypsilanti, Jennifer A. Akiyama, Brandon J. Mannion, Ingrid Plajzer-Frick, Catherine S. Novak, Anne N. Harrington, et al. 2021. “Ultraconserved Enhancer Function Does Not Require Perfect Sequence Conservation.” Nature Genetics 53 (4): 521–28.

Srivastava, D., and E. N. Olson. 1997. “Knowing in Your Heart What’s Right.” Trends in Cell Biology 7 (11): 447–53.

Stennard, Fiona A., Mauro W. Costa, David A. Elliott, Scott Rankin, Saskia J. P. Haast, Donna Lai, Lachlan P. A. McDonald, et al. 2003. “Cardiac T-Box Factor Tbx20 Directly Interacts with Nkx2-5, GATA4, and GATA5 in Regulation of Gene Expression in the Developing Heart.” Developmental Biology 262 (2): 206–24.

Taher, Leila, David M. McGaughey, Samantha Maragh, Ivy Aneas, Seneca L. Bessling, Webb Miller, Marcelo A. Nobrega, Andrew S. McCallion, and Ivan Ovcharenko. 2011. “Genome-Wide Identification of Conserved Regulatory Function in Diverged Sequences.” Genome Research 21 (7): 1139–49.

Taskiran, Ibrahim I., Katina I. Spanier, Hannah Dickmänken, Niklas Kempynck, Alexandra Pančíková, Eren Can Ekşi, Gert Hulselmans, et al. 2023. “Cell-Type-Directed Design of Synthetic Enhancers.” Nature, December, 1–9.

Torchiano, Marco. 2016. Effsize - a Package for Efficient Effect Size Computation. Zenodo. 10.5281/ZENODO.1480624.

Villar, Diego, Camille Berthelot, Sarah Aldridge, Tim F. Rayner, Margus Lukk, Miguel Pignatelli, Thomas J. Park, et al. 2015. “Enhancer Evolution across 20 Mammalian Species.” Cell 160 (3): 554–66.

Vinga, Susana. 2014. “Information Theory Applications for Biological Sequence Analysis.” Briefings in Bioinformatics 15 (3): 376–89.

Visel, Axel, Matthew J. Blow, Zirong Li, Tao Zhang, Jennifer A. Akiyama, Amy Holt, Ingrid Plajzer-Frick, et al. 2009. “ChIP-Seq Accurately Predicts Tissue-Specific Activity of Enhancers.” Nature 457 (7231): 854–58.

Visel, Axel, James Bristow, and Len A. Pennacchio. 2007. “Enhancer Identification through Comparative Genomics.” Seminars in Cell & Developmental Biology 18 (1): 140–52.

Wu, Tianzhi, Erqiang Hu, Shuangbin Xu, Meijun Chen, Pingfan Guo, Zehan Dai, Tingze Feng, et al. 2021. “ClusterProfiler 4.0: A Universal Enrichment Tool for Interpreting Omics Data.” Innovation (Cambridge (Mass.)) 2 (3): 100141.

Xiao, Feng, Xiaoran Zhang, Sarah U. Morton, Seong Won Kim, Youfei Fan, Joshua M. Gorham, Huan Zhang, et al. 2024. “Functional Dissection of Human Cardiac Enhancers and Noncoding de Novo Variants in Congenital Heart Disease.” Nature Genetics 56 (3): 420–30.

Zaidi, Samir, Murim Choi, Hiroko Wakimoto, Lijiang Ma, Jianming Jiang, John D. Overton, Angela Romano-Adesman, et al. 2013. “De Novo Mutations in Histone-Modifying Genes in Congenital Heart Disease.” Nature 498 (7453): 220–23.

Zhang, Xiaoyu, Irene M. Kaplow, Morgan Wirthlin, Tae Yoon Park, and Andreas R. Pfenning. 2020. “HALPER Facilitates the Identification of Regulatory Element Orthologs across Species.” Bioinformatics (Oxford, England) 36 (15): 4339–40.

Zielezinski, Andrzej, Hani Z. Girgis, Guillaume Bernard, Chris-Andre Leimeister, Kujin Tang, Thomas Dencker, Anna Katharina Lau, et al. 2019. “Benchmarking of Alignment-Free Sequence Comparison Methods.” Genome Biology 20 (1): 144.

Zielezinski, Andrzej, Susana Vinga, Jonas Almeida, and Wojciech M. Karlowski. 2017. “Alignment-Free Sequence Comparison: Benefits, Applications, and Tools.” Genome Biology 18 (1): 186.

